# Tracking HIV-1 DNA fate from Cell Culture to Humanized mice Tissues

**DOI:** 10.1101/2025.11.25.690387

**Authors:** Jean-Sebastien Diana, Evgeny Tatirovsky, Chiara Tomasini, Celine Cuche, Selen Ay, Viviana Scoca, Marina Cavazzana, James P. Di Santo, Francesca Di Nunzio

**Author notes:** Shared first authors (Jean-Sebastien Diana, Evgeny Tatirovsky). Shared last authors (James P. Di Santo, Francesca Di Nunzio).

## Abstract

Unambiguous identification of HIV reservoirs is essential for their characterization and for developing curative strategies. A major obstacle to curing HIV is the persistence of silent viral genomes in host cells that evade immune detection and resist therapy. Here, we present a fluorescence microscopy-based method for direct in vivo visualization of full-length double-stranded HIV-1 DNA, an achievement not previously realized. To achieve this, we developed an advanced imaging platform adapted for replication-competent HIV-1. This system leverages a bacterial-derived two-component tagging strategy, where a fluorescent protein (OR-GFP), efficiently expressed in xenografted human immune cells, specifically binds to an engineered ANCH3 tag sequence integrated into the viral genome.

The tagged virus infects CD4+ T cells both *in vitro* and *in vivo*, producing bright nuclear puncta corresponding to individual viral genomes. Multiple nucleation sites within ANCH3 enable a stable, shortened tag form for persistent and long-term tracking. Transcriptional profiling during acute infection revealed both transcriptionally active and silent genomes in spleen, lymph nodes, and bone marrow, with silent forms enriched in lymph nodes and marrow, supporting early reservoir establishment. This live-cell imaging strategy enables high-specificity detection of latent HIV, offering a transformative tool for studying reservoir dynamics and guiding future cure strategies.

## Introduction

The HIV-1 replication cycle has been extensively studied, yielding valuable insights into the critical interactions that occur between the virus and host factors during permissive infection (1). This knowledge has been instrumental in the development of antiretroviral drugs (ART) that target different stages of the virus life cycle (2). While ART is effective in inhibiting HIV-1 replication, it falls short of eradicating the infection (3–11). This is primarily due to the establishment of persistent infection reservoirs, which remain largely untouched by ART and represent a major barrier to achieving an HIV cure (12, 13).

The enduring presence of reservoirs stems from the stable integration of viral DNA into the host cell’s genetic material, facilitating persistent infection throughout the cell’s lifespan and its descendants (14, 15). Remarkable strides have been made in understanding the virological attributes of HIV reservoirs, including the integrity of proviruses, their potential for reactivation, and the landscape of integration sites (16, 17). Recently, single-cell analytical techniques have been used to investigate the proteomic, epigenetic, transcriptomic, and phenotypic profiles of individual HIV-infected cells, including memory CD4+ T cells from both peripheral blood and lymph nodes of individuals living with HIV-1 (18–20). Nonetheless, despite these significant advances in the characterization and indirect quantification, the cellular identity of the HIV reservoir *in vivo* still remains enigmatic.

Recent advances in imaging technologies have provided new tools for studying the HIV-1 life cycle and reservoir formation (21–24). Combination of imaging microscopy with fluorescently labelled viral components enables the examination of HIV-1 biology at the single-cell level (25–30). However, the generation of fluorescently-modified HIV-1 genomes has been shown to impact the viral replication fitness (23). In addition, the high mutation rate of HIV-1 frequently results in loss of the engineered fluorescent tags (24). As such, creating a replication-competent HIV-1 with a stable tracking modification suitable for *in vivo* reservoir detection (for example in HIV-1-infected humanized mice) remains a challenge.

In this study, we have engineered a full-length HIV-1 genome to accommodate the ANCHOR^TM^ system (26, 31) therby allowing detection of HIV-1 DNA in cells expressing the specific OR-GFP protein. The ANCHOR^TM^ system is a bipartite labeling method derived from the bacterial *parABS* chromosome segregation machinery, comprising a DNA sequence (ANCH3) that binds a specific protein (OR). Previous studies have used ANCH3/OR interactions to track viral DNA from non-replication-competent viruses as microscopically visible fluorescent nuclear punctae (26, 32). Here we describe a novel ANCH3-modified HIV-1 virus that can effectively infect cell lines and primary human cells *in vitro*, facilitating the identification of cells harbouring viral DNA through confocal microscopy. Using an *in vivo* human immune system (HIS) mouse model (33), we found that the ANCH-modified HIV-1 robustly replicated during the acute infection period. Interestingly, we observed adaptation of the ANCH tag during viral replication *in vitro* and *in vivo*, resulting in selection of shorter form containing four nucleation sites, which allowed decoration of OR-GFP along the surrounding HIV-1 dsDNA genomic sequence. Such HIV-1 dsDNA was visible as bright spot easily detectable by light microscopy in primary lymphocytes or in CD4+ T cells isolated from different organs of ANCH-HIV-1-infected HIS mice. When combined with the RNA-FISH technique, this ANCHOR^TM^ method also enabled the differentiation between non-transcribing and actively transcribing HIV-1 DNA, both *in vitro* and *ex vivo*. Lastly, we successfully detected HIV-1 DNA in live cells using spinning disk microscopy. In summary, our approach offers an effective means of investigating mechanisms related to HIV-1 infection and formation of latent reservoirs *in vivo*.

## Results

### An ANCH3-modified replication-competent HIV-1 for the live tracking of the viral genome

Here, we aim to adapt ANCHOR^TM^ system to a genetically modified, replication-competent HIV-1 strain to enable live tracking of the viral genome during the complete replication cycle. We first set out to generate a NL4.3 AD8 HIV-1 replication-competent ANCHOR^TM^ constructs (Figure 1A). The functional ANCH3 sequence has been engineered to be 400bp or 1kb long, and contains several nucleation sites for binding the OR proteins. ANCH3 sequence has been inserted at the place of the iresGFP between the Nef gene and the 3’LTR region (Figure 1A). OR-fluorescent protein (OR-GFP) specifically recognizes the double stranded ANCH3 sequence, which is generated at the end of the reverse transcription process. Thus, bright nuclear spots should label intact double stranded vDNA (31, 32). The substantial accumulation of OR-GFP protein onto its cognate ANCH3 sequence (up to 500 OR-GFP per single ANCH3 sequence) is enough to detect single DNA molecules (31). To further enhance the adaptability of the ANCHOR^TM^ constructs within the human primary cells, we replaced the previously used CMV promoter (26, 32) with the EF1alpha (EF1a) promoter in the transfer vector carrying OR-GFP (Figure S1A). We generated two lentiviral vectors containing either the long or short version of the EF1α promoter. Since both showed similar transduction efficiency, we proceeded with the long EF1α promoter (Figure S1A).

**Figure 1.**
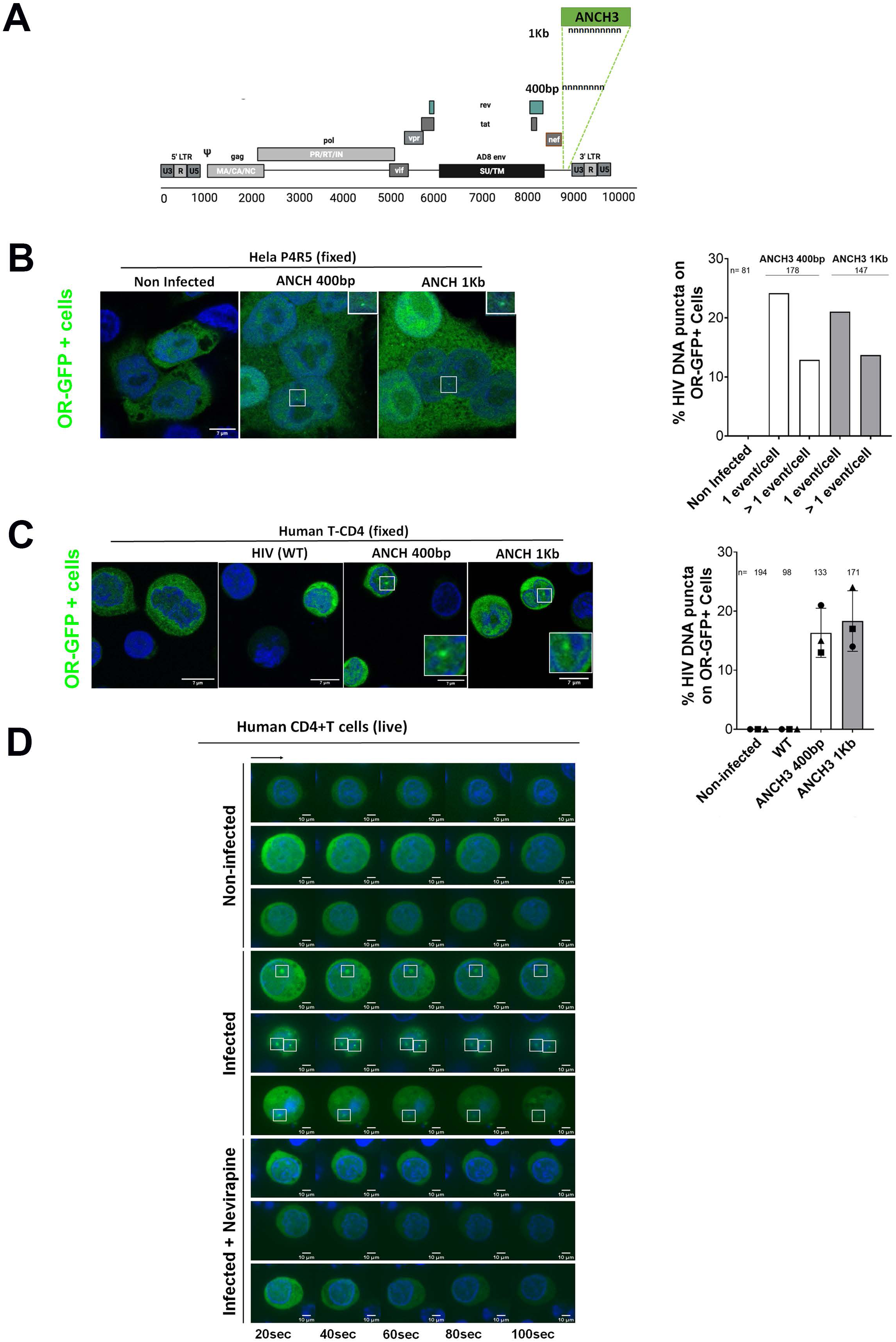
*In vitro* infectivity and DNA spot detection of a replication competent HIV-1 carrying 1 Kb and 400bp ANCH3 sequence. **A,** schema of HIV-1 NL4.3 AD8 wild type (WT), NL4.3 AD8 ANCH3 1kb or NL4.3 AD8 ANCH3 400bp. **B,** detection of nuclear HIV-1 DNA forms by confocal microscopy and labeling of vDNA in HeLa P4R5 following infection with NL4.3 AD8 ANCH3 400bp and NL4.3 AD8 ANCH3 1Kb (MOI 50, 30 hours p.i.). The nucleus is labeled by hoechst in blue and the vDNA via OR-GFP in green (OR-GFP protein has not the NLS so it is mainly localized in the cytoplasm, inside the nucleus OR-GFP forms bright spots in association with the vDNA). Quantification of the % of cells carrying vDNA spot events summarized in the graph on the right. **C**, detection of nuclear HIV-1 DNA forms from NL4.3 AD8 ANCH3 400bp and NL4.3 AD8 ANCH3 1Kb in activated CD4+T cells (MOI 20, 3 days p.i.) by confocal microscopy. Labeling are as in B. The graph in the right shows the percentage of HIV DNA signal on the total of OR-GFP positive cells. **D**, *in vitro* live-tracking of the viral DNA (ANCHOR) in primary CD4+ T cells seeded on collagen and imaged using a spinning disk microscope. Images were acquired continuously (20sec per frame). Cells imaged were infected with or without NEV (10µM) or uninfected. Nuclei are in blue, OR-GFP protein is in green, vDNA is shown by intranuclear green spots of OR-GFP proteins.

Both HIV-1 NL4.3 AD8 ANCH3 400bp and ANCH3 1 Kb were able to infect *in vitro* OR-GFP transduced CCR5+ cell lines (Hela P4R5) and primary CD4+ activated T cells (Figures 1B,C and S1A,B). Indeed, the reverse transcribed HIV-1 genome was detected in both conditions of NL4.3 AD8 ANCH3 1kb and ANCH3 400bp infection in HeLa P4R5 cells and CD4+T cells isolated from various healthy donors (Figures 1B, C). We observed that the shorter tag form, the 400 bp ANCH3 variant, can be used to live-track viral DNA in infected primary CD4+ T cells using spinning disk confocal microscopy (Figure 1D, Movies 1,2,3). Interestingly, the 400bp ANCH3 variant exhibits a sensitivity comparable to that of the 1Kb version in detecting HIV proviral DNA (Figures 1B,1C and S1A). This observation suggests that the sensitivity is not necessarily dependent on the size of the ANCH3 tag or the number of nucleation sites, since the OR proteins first bind to the specific sequence and then spread to the surrounding chromatin regions (31).

### *In vitro* and *in vivo* fitness of ANCH3-modified HIV-1 viral strains

Modification of HIV-1 genomes with fluorescent tags frequently results in a loss of viral fitness (34). We therefore tested the replicative capacity and transcriptional activity of ANCH3-modified HIV-1 strains. Modified-HIV-1 carrying different ANCH3 insert size generate similar viral titers (4×10^9^ TU/mL). Next, we assessed *in vitro* viral fitness by measuring p24 production at different days after CD4+T cells infection, comparing WT and ANCH3-tagged HIV-1 NL4.3. Interestingly, we found that the NL4.3 AD8 ANCH3 1kb was unable to replicate *in vitro* (Figure 2A). When analyzing the ratio between the number of physical particles (PP) and Transducing Unit (TU) values, the viral production quality of the virus carrying the 1Kb sequence was found to be 37-fold lower than that of the ANCH3 400bp tagged virus. Intriguingly, the latter exhibited a comparable functional viruses value to WT HIV-1 (PP/TU= 73 vs. 63, respectively). These findings suggest that the longer ANCH3 form might exert a negative influence on viral RNA genome packaging and on the formation of viral particles, both of which are essential for achieving optimal viral fitness. In contrast, the HIV NL4.3 AD8 ANCH3 (400bp) strain demonstrated a fitness level similar to that of the WT virus *in vitro* (Figures 2A,B). We further tested the viral fitness of NL4.3AD8 ANCH3 400bp and ANCH3 1kb *in vivo* using a human immune system (HIS) mouse model (35). We assessed HIV-1 replication in 12-week-old BRGS A2DR2 HIS mice (see materials and methods for details) after injection of 10^5^ TCID50 of WT and HIV-1 WT-ANCHOR viruses, carrying ANCH3 1Kb or 400bp (Figure 2C). Consistent with the *in vitro* observations, we found no replication of NL 4.3 AD8 HIV ANCH3 1Kb in BRGS A2DR2 HIS mice. However, NL 4.3 AD8 HIV ANCH3 400bp infection resulted in robust plasma viremia, similarly to the untagged virus (Figure 2D). Next, we evaluated the persistence of the tag ANCH3 in infected HIS mice until 21 days post-infection. We performed *in vitro* reverse transcription on viral RNA extracted from the plasma of HIS mice, followed by a two-step nested PCR using primers that anneal to the viral genome at the ends of the cloned tag. This allowed us to amplify the ANCH3 sequence inserted into the full-length HIV-1 genome three weeks post-infection. Gel analysis consistently revealed a shorter band of approximately 250 bp, instead of the expected 400 bp, in all mice tested (Figure 2E). Sequencing of this band showed the presence of 4 out of 8 nucleate sites. Interestingly, the missing sequence was located between two palindromic SmaI restriction sites.

**Figure 2.**
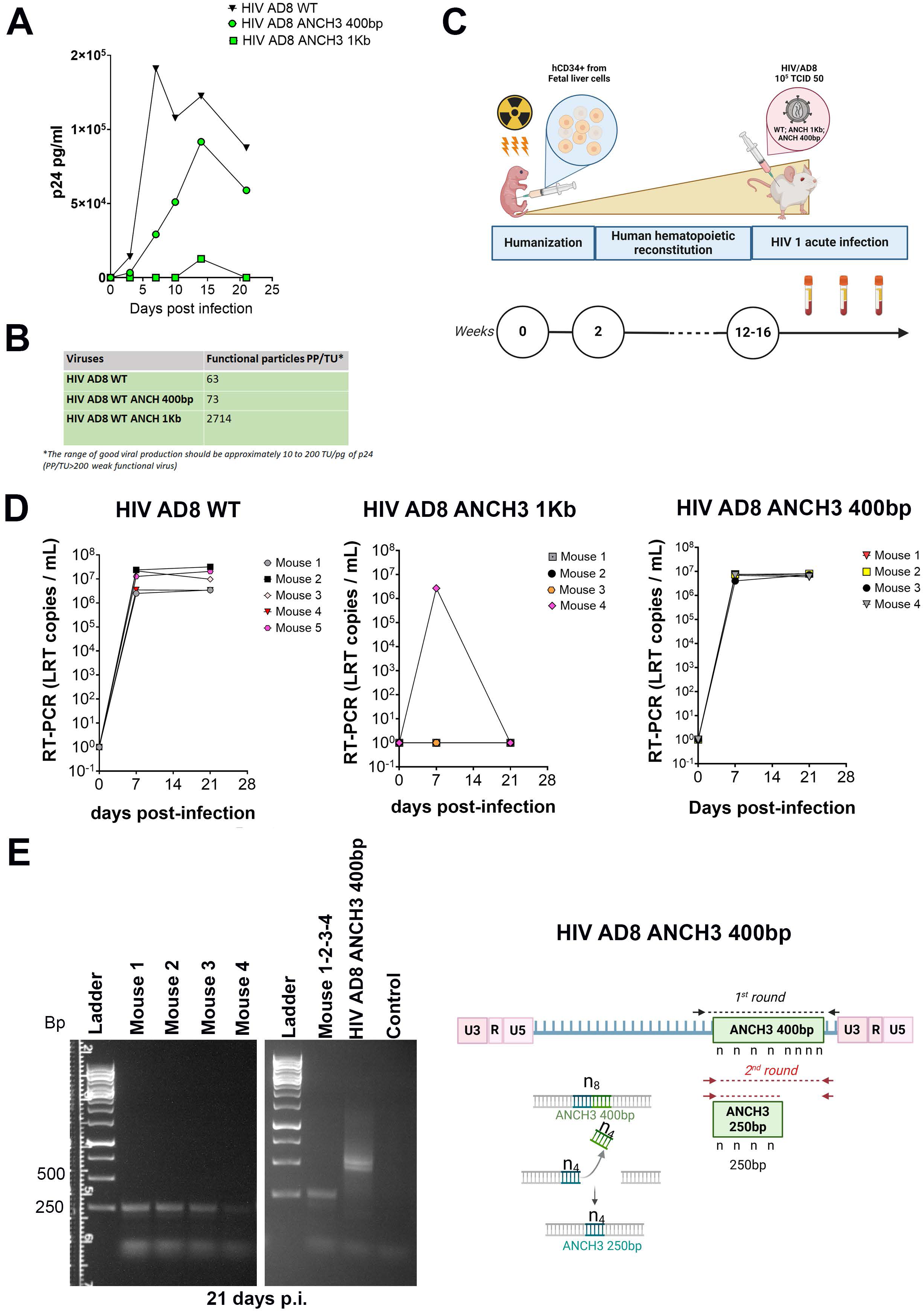
Viral fitness of NL4.3AD8 ANCH3 400bp and ANCH3 1 kb in BRGS A2DR2 fetal liver humanized mice. **A**, viral fitness *in vitro* by p24 assay (0, 3, 7, 10, 14, 21 days p.i.) HIV NL4.3 AD8 WT, NL4.3 AD8 ANCH3 400bp and NL4.3 AD8 ANCH3 1Kb. **B**, the table reports the values of functional particles calculated as ratio between physical particles and transducing units (PP/TU) for the three viruses tested. **C**, experimental schema and timing of NL4.3 AD8 HIV-1 WT or genetically modified with ANCH3 (1Kb and 400bp) infection in BRGS A2DR2 HIS mice. BRGS A2DR2 HIS mice: newborn (3-to 5-days-old) pups received sublethal irradiation (3 Gy) and 5 × 10^4^ CD34+CD38– human fetal liver cells or 2 × 10^5^ to 4 × 10^5^ CD34+ human cord-blood were injected intrahepatically. Once the engraftment of mice was confirmed, twelve-weeks-old reconstituted HIS mice were inoculated with 10^5^ TCID50 (tissue culture infective dose required for 50% infection) of NL4.3 AD8 HIV-1 ANCH3 1Kb and 400 bp. **D,** Infected mice underwent to plasma viremia test at the indicated times in the acute phase (d0, d7, and d21). HIV-1 general viral load quantified by late reverse transcripts (LRTs) qPCR from vRNA extracted from plasma. **E**, On the left, amplified ANCH3 sequence in viral RNA extracted from mice plasma samples infected by HIV NL4.3 AD8 ANCH3 400 bp for 21 days. The viral RNA isolated was firstly reverse transcribed. The cDNA was then amplified using specific primers for the HIV sequences flanking ANCH3 sequence and vDNA genome followed by a nested PCR. Amplicons from the nested PCR of each mouse samples are run on agarose gel 1% as shown in the panel of the gel on the left. Panel on the right shows amplicons from a mix of samples of four infected mice and the next lane is loaded with he amplicon derived from the positive control (virus) (the NL4.3 AD8 ANCH3 400 used to infect the four mice in D and E). On the right, cartoon of the PCR strategy (created with BioRender).

### *In vitro* and *in vivo* selection of an optimal ANCH3 Tag in replicative HIV-1

To investigate whether the ANCH3 tag was modified during viral replication, we infected CD4+ T cells derived from two healthy donors and sequentially harvested viral supernatants to infect new cells over five consecutive cycles (Figure 3A). RT-PCR amplification of the ANCH3 tag sequence revealed a 400 bp band at the time of infection and up to 3 days post-infection (dpi). At 7 dpi a shorter band began to appear which emerged as the only detectable and stable band observed in the last time point measured in our experiment, 21 dpi (Figure 3B). Sequencing of the ANCH3 amplicon at 21 dpi confirmed the presence of four nucleation sites for the OR-binding protein.

**Figure 3.**
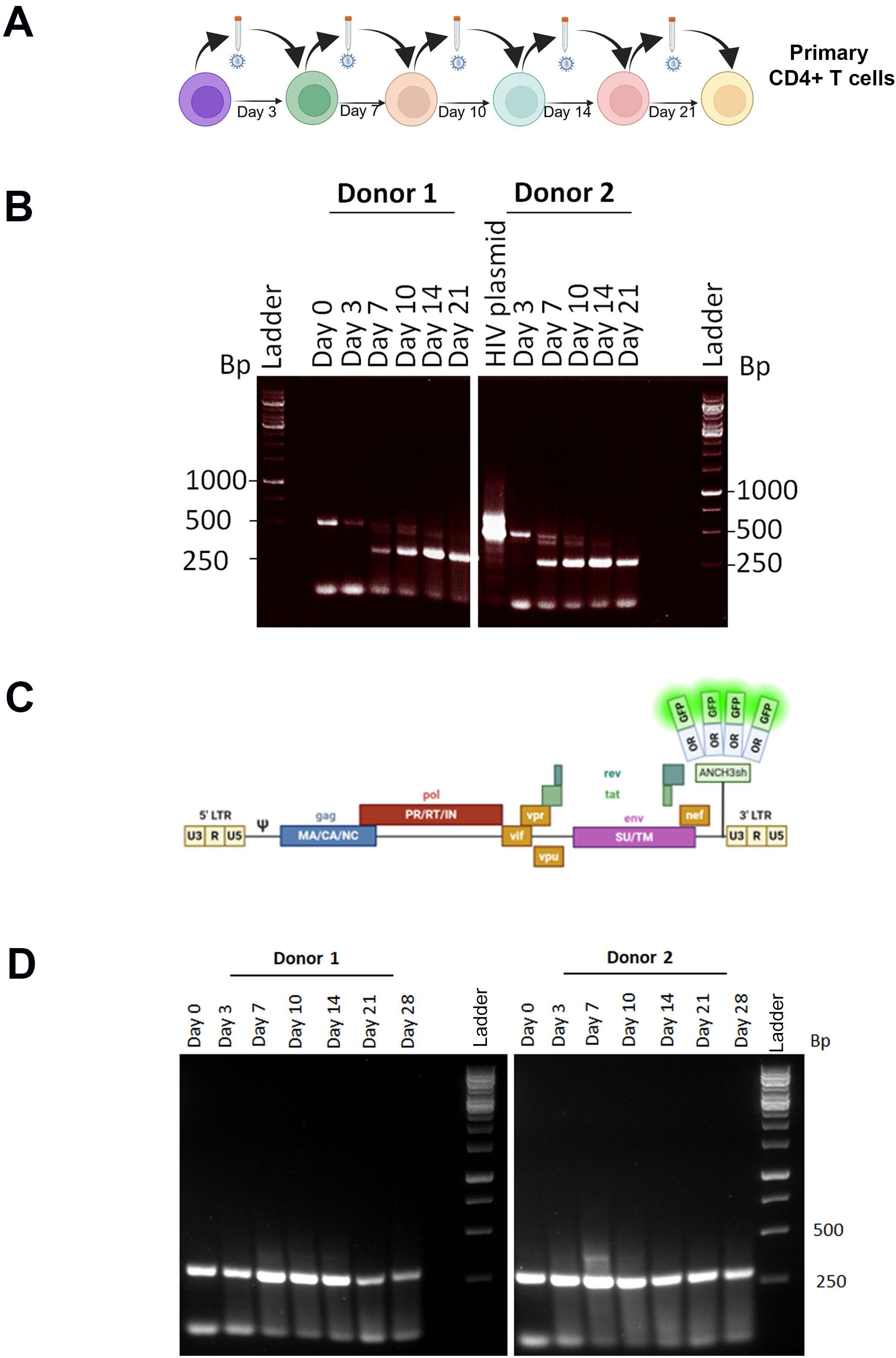
Evaluation of the stability of the ANCH3 sequence over multiple passages of infection in CD4+ T cells from healthy donors. **A**, schema of multiple passages of infection in primary CD4+ T cells (created with BioRender). **B,** amplicons of ANCH3 sequence in viral RNA extracted from the supernatant of Primary CD4+ T cells of two donors (Donor 1 and Donor 2) infected by HIV NL4.3 AD8 ANCH3 400 bp, following the model exposed in Figure 2E. The amplified sequence was run on agarose gel 1.3%. **C,** cartoon illustration of the new virus carrying the naturally selected 250 bp sequence at 21 days post-infection, based on its sequence (created with BioRender). **D,** Infection of primary T cells from donors has made according to the schema illustrated in panel A using the HIV-1 NL4.3 AD8 ANCH3sh (250 bp) described in panel C. Amplicons of ANCH3 sequence of viral RNA extracted were amplified in a single round using external primers at the sequence ANCH3. The procedure for reverse transcription, PCR amplification and electrophoresis on agarose gel followed the same steps shown in panel B.

Based on the sequence obtained from the adapted virus (*in vitro* and *in vivo*), we generated a new viral construct carrying the short form (250bp) of the ANCH3 tag obtained by sequencing (Figure 3C). We then infected CD4+ T cells from two healthy donors and collected supernatants over time to perform RT-PCR across different viral passages, from day 0 to day 28. We observed a single, stable band throughout all analyzed passages, indicating the 250bp ANCH3 tag was stable and not further modified after multiple rounds of infection in primary CD4+ T cells (Figure 3D).

### Tracking HIV-1 DNA *in vivo* using a humanized mouse model

Our aim was to establish a proof of concept that would pave the way for future applications of HIV-1 WT-ANCHOR system using an appropriate animal model, allowing us to delve into the investigation of reservoir cells and secondary lymphoid tissues. It has been previously published that HIS mice overexpressing mouse TSLP (BRGST HIS mice) generate a full complement of secondary lymphoid tissues (lymph nodes and spleen) and primary lymphoid tissues as the bone marrow that harbor human T cells susceptible to HIV-1 infection (35). Thus, we infected fetal liver (FL) CD34+ engrafted BRGS A2DR2 mice with HIV carrying the ANCH3 sequence of 400bp and three weeks post-infection we isolated total PBMC and positive and negative fractions of splenic CD4+ cells. We amplified ANCH3 tag (primers and probe amplify ∼160nt at the 5’ of ANCH3 tag) from those spleen cells quantifying the ANCH3 sequence at a rate of 0.2 copies per CD4+ T cells (Figure 4A). We subsequently transduced CD4+ T cells isolated from the spleen of HIV-infected HIS mice with a lentiviral vector encoding OR-GFP under the control of the EF1α promoter. Confocal microscopy revealed distinct and bright GFP puncta within the nuclei of infected cells (Figure 4B). These results collectively demonstrate the feasibility of tracking the final products of viral reverse transcription in the nucleus using an *in vivo* model.

**Figure 4.**
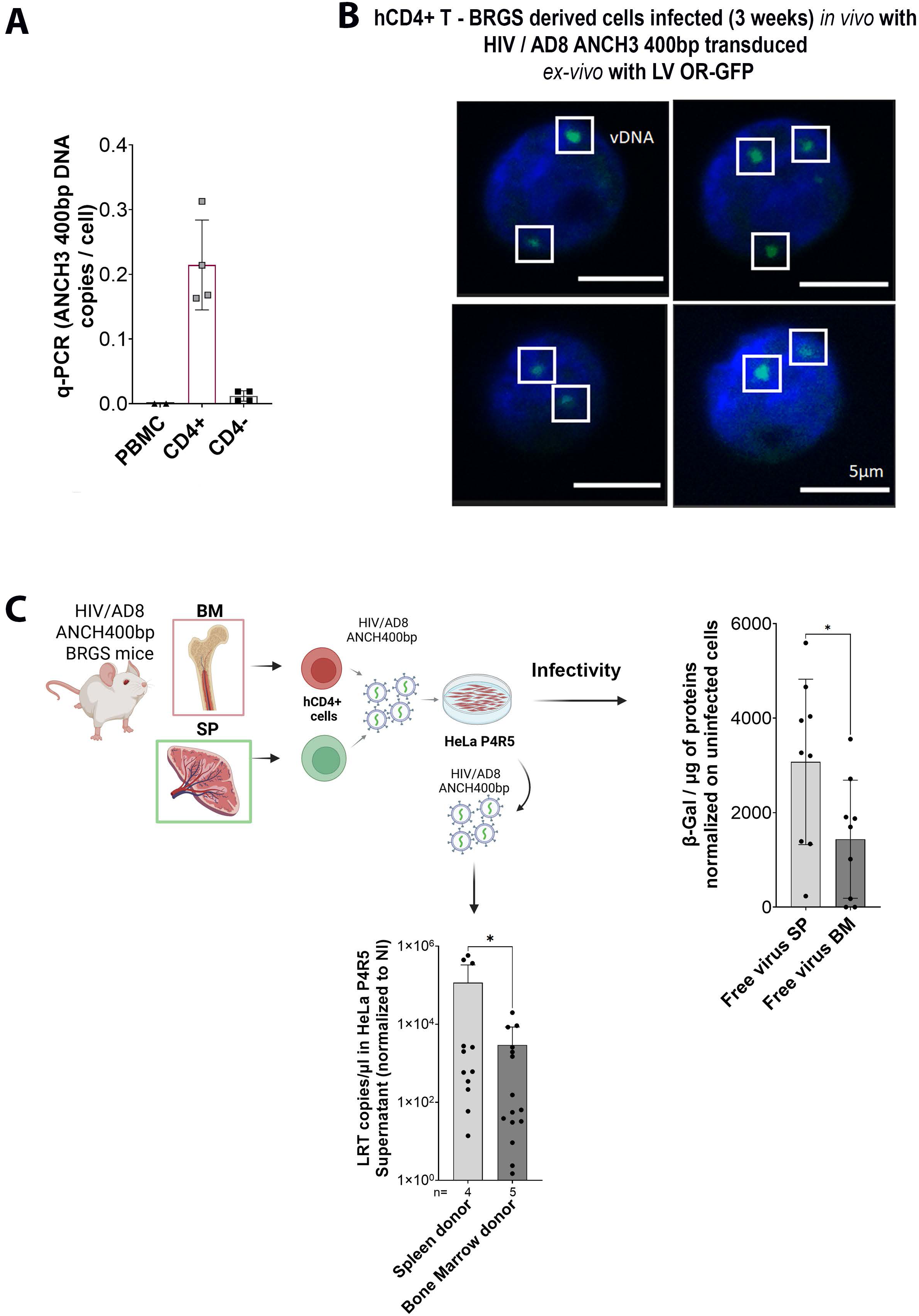
HIV carrying ANCH3 400 bp allows viral DNA visualization in the nucleus of cells isolated from HIS mice, and these cells can spread the virus under *in vitro* conditions. **A,** ANCH 400bp DNA qPCR quantification in sorted CD4+ and CD4-splenocytes and in PBMC (d.p.i 21). **B**, vDNA HIV spot detection in CD4+ sorted and *ex vivo* OR-GFP transduced splenocytes from infected mice. **C,** CD4⁺ T cells were isolated from spleen (SP) and bone marrow (BM) of HIS mice infected with HIV-1 NL4.3 AD8 carrying the ANCH3 400 bp and activated *in vitro* for 3 days. The resulting supernatant was then collected and used to infect HeLa P4R5 cells. Infectivity in HeLa P4R5 was assessed using a β-galactosidase assay (right panel), while viral production from HeLa P4R5 cells was measured by qRT-PCR using primers specific to the viral genome in the supernatant (bottom graph). Statistical test: T test, p values are shown in the graph.

Next, we aimed to demonstrate the capacity of the ANCH3-tagged virus to spread *in vivo*. To this end, we used BRGS A2DR2 HIS mice for subsequent experiments (Figure 4C). CD4⁺ T cells were isolated from the bone marrow and spleen three weeks post-infection, then activated *in vitro* with cytokines for three days. The supernatants from these activated CD4⁺ cells were collected and used to infect HeLa P4R5 cells over a three-day period. Infectivity assays based on β-galactosidase activity confirmed that HeLa P4R5 cells were successfully infected by free virus present in the supernatants derived from both spleen and bone marrow CD4⁺ T cells. On average, HeLa P4R5 cells were twice as infected by supernatants from spleen-derived cells compared to those from bone marrow. Consistently, RT-qPCR analysis of the supernatants from HeLa P4R5 cells previously exposed to spleen- or bone marrow-derived cell supernatants revealed significantly higher levels of viral RNA in cells treated with spleen-derived supernatant compared to those treated with bone marrow-derived supernatant (Figure 4C). Taken together, these results indicate that spleen CD4⁺ T cells produce more virus than bone marrow cells in this controlled *in vitro* assay using *in vivo* samples.

We found that human CD34+ hematopoietic precursors derived from cord blood (CB) were more easily genetically modified using lentiviral vectors compared to similar cells derived from human fetal liver cells (Figures S2A,B,C,D). Both human fetal liver-derived and human cord blood-derived CD34+ cells have been previously shown to efficiently reconstitute HIS mouse recipient strains (36–39). As such, we employed human cord blood-derived CD34+ cells for OR(3)-GFP transduction and generation of BRGST A2DR2 ^OR-GFP^ HIS mice (Figure 5A). Transduced CB hCD34+ cells (∼77.2% OR-GFP+ cells) were injected into irradiated BRGS/T pups (Figures 5A,B), and, after at least 10 weeks, the mice were validated for immune reconstitution and OR-GFP expression. We found that the humanization, defined as a percentage of human CD45+ cells from the total CD45+ (mouse and human) cells, in CB-engrafted BRGS/T A2DR2^OR-GFP^ did not differ significantly from the FL-engrafted BRGS/T A2DR2 mice: 28% (range (8-42%) vs. 26% (range (10-60%) (Figure 5C). The percentage of humanization has been calculated also in individual organs: spleen, bone marrow and lymph nodes (Figure 5D). Reconstitution of BRGST A2DR2 and BRGS/T A2DR2^OR-GFP^ generated the same multilineage of hCD45+ cells composed by human T cell population (hCD45+ CD3+ CD4+ and CD8+), human B cell (CD19+), human myeloid cells (CD3-CD19-HLA-DR+) and NK (NKp46+ CD94+) (Figure S3). Results for humanization and OR-GFP+ cells engraftment are summarised in supplementary data (Figures S3,S4,S5,S6,S7) Altogether, the engraftment, was confirmed and sufficient for HIV-1 replication in 15 out of 17 BRGS/T A2DR2^OR-GFP^ (Table S1, Figure S3). We assessed OR-GFP expression in human hematopoietic cells from bone marrow, spleen, and lymph nodes in BRGS/T A2DR2 HIS mice at ∼ sixteen weeks post-engraftment and three weeks post-infection. We used confocal microscopy on CD4+ sorted cells from organs to detect OR-GFP protein expressing cells (Figure 5E). Confocal microscopy confirmed that CD4+ OR-GFP+ cells were detectable in bone marrow, spleen (∼70%) and in the lymph nodes (∼45%)(Figure 5E). In addition, the expression of OR-GFP remained stable over time as indicated from the percentage of change in the numbers of OR-GFP+ hCD45+ cells and OR-GFP+ CD4+ T cells in the blood over several weeks post engraftment (Figures 5F,G). In conclusion, OR-GFP transduction and engraftment of human hematopoietic precursors allowed for normal generation of HIS mice with subsequent potential for HIV infection. We next analyzed the HIV-1 NL4.3 AD8 ANCH3 400bp fitness in BRGS/T A2DR2^OR-GFP^ HIS mice. BRGS/T A2DR2^OR-GFP^ mice challenged with HIV-1 NL4.3 AD8 ANCH3 400bp showed robust viremia up to ∼10^7^ copies/ml (Figure 6A). We again documented stability of the ANCH3 400bp tag sequence throughout the infection in BRGS/T A2DR2^OR-GFP^ HIS mice (Figure 6B). Acute infection with ANCH3-modified HIV-1 led to a progressive rise in viremia and a corresponding decline in CD4+ T cells, mimicking the acute phase of infection observed with WT HIV-1 (Figure 6B). The persistence of OR-GFP⁺ cells was also evaluated up to three weeks post-infection.The changes in the percentage of OR-GFP⁺ CD4⁺ cells among human CD45⁺ cells (Figure 6C) and in their numbers in the blood (Figure 6D) indicate stable OR-GFP expression in the context of HIV-1 infection.

**Figure 5.**
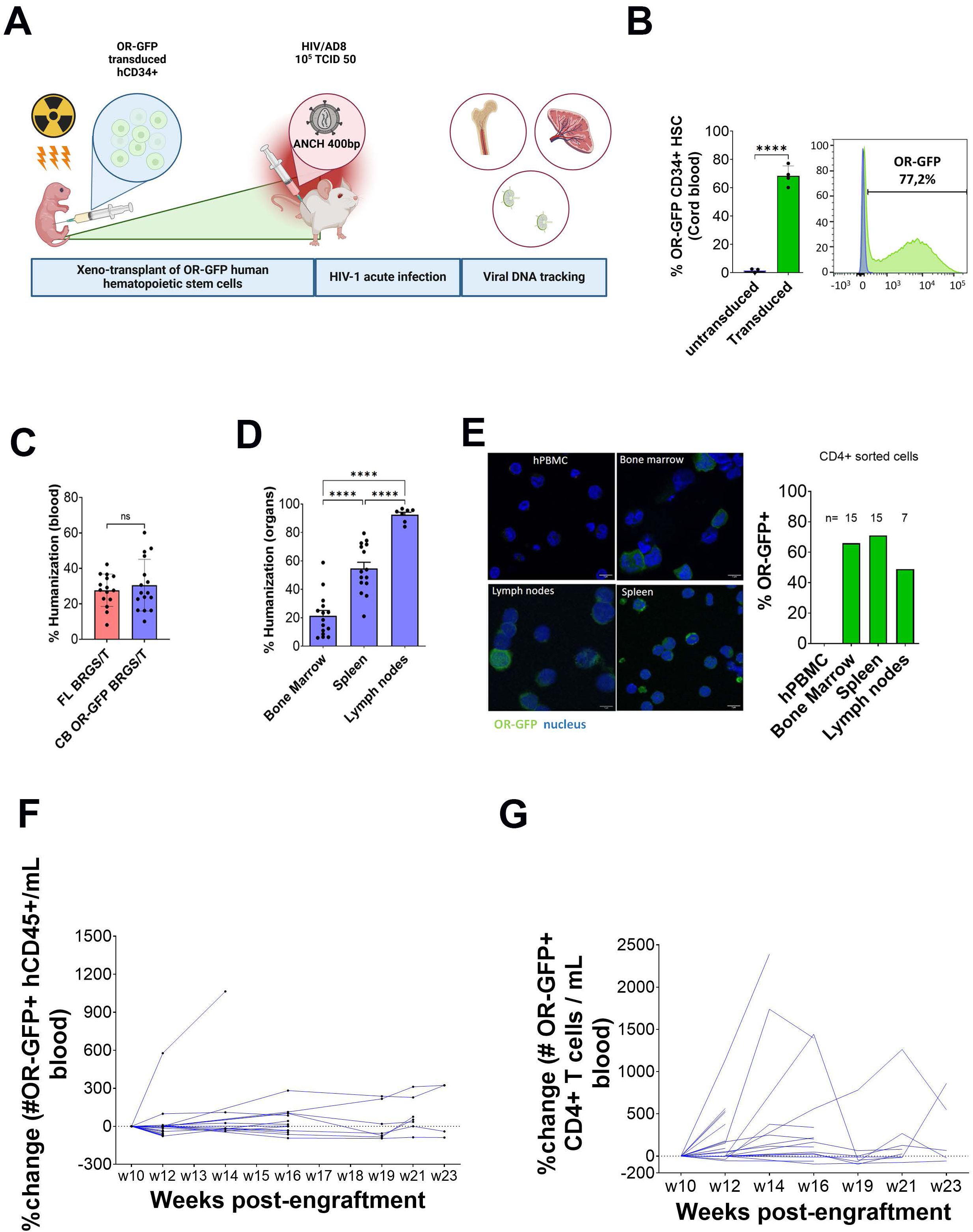
BRGS/BRGST A2DR2 OR-GFP xenografted model: *in vivo* model of HIV-1 DNA detection. **A**, experimental schema and timing of BRGS/BRGST A2DR2^OR-GFP^ (BALB/c RAG2^-/-^ IL2Rg^-/-^ SirpaNOD2 ^-/-^ A2DR2). hCD34+ cells were pre-activated overnight and OR(3)-GFP transduced. The transduction medium was X-vivo 20 supplemented with SCF, Flt3-L, TPO and IL3. We used non-toxic enhancers of transduction: Protamine sulfate, PGE2, and culture plates were coated with fibronectin. Newborn (3-to 5-days-old) pups received sublethal irradiation (3 Gy) and 2 × 10^5^ to 4 × 10^5^ OR(3)-GFP transduced CD34+ were injected intrahepatically (created with BioRender). **B**, hCD34+ OR-GFP transduction efficacy for BRGS/T A2DR2 xenograft model quantified by the OR-GFP+ expression in hCD34+ cells by FACS after 7 days of hCD34+ transduced culture. **C**, primary engraftment, defined by the FACS detection of hCD45+ cells as a fraction of total CD45+ cells. The humanization was calculated (hcd45*100/(m+hcd45)) for PBMC and in cell suspension derived from bone marrow, spleen and lymph nodes (d.p.i 21). **D**, Percentage of OR-GFP + CD4+ cells sorted from different organs from the xenografted HIS mice measured by FACS. **E**, Left figure shows the representative confocal images with nuclei in blue and OR-GFP in green from bone marrow, spleen and lymph nodes. Right panel – quantification of the %OR-GFP + cells in the organs and PBMCs. **F**, The percentage of change in the numbers of OR-GFP+ total hCD45+ cells and **G**, CD4+ T cells per mL, measured by flow cytometry in peripheral blood and calculated using initial values as a baseline.

**Figure 6.**
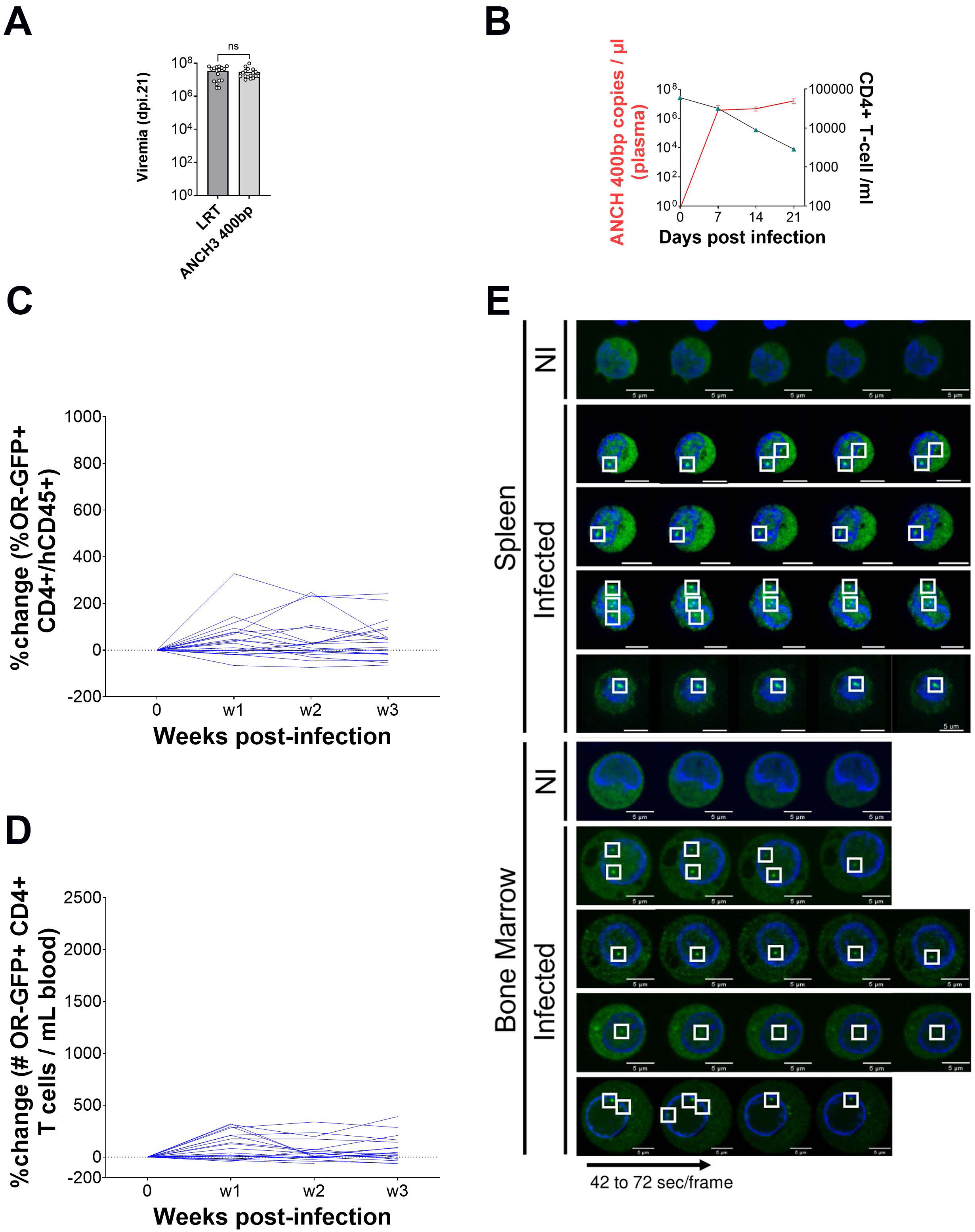
Fitness and Live Tracking of the HIV Genome in Humanized Mice. **A,** RT-PCR was performed on viruses circulating in the blood at 3 weeks post-infection using two sets of primers: one set targeting late reverse transcripts spanning the U5–gag junction, and another set specific for the 5′ ANCH3 sequence with respective probes. **B**, specific viremia in plasma on dpi 7,14 and 21 by RT qPCR with ANCH3 400bp (red), merged with the CD4+ T cells count depletion (black) during the acute phase of infection in seven BRGS A2DR2 ^OR-GFP^ mice. **C**, The percentage of change in the percentage of OR-GFP+ CD4+ from total hCD45+ cells and **D**, in their numbers per mL, measured by flow cytometry in peripheral blood and calculated using initial values as a baseline. **E**, *in vivo* live tracking of the viral DNA (ANCHOR) in CD4+ T cells isolated from spleen and bone marrow of BRGS A2DR2 ^OR-GFP^ mice, plated after a CD4+T sorting in a polymer-coverslip bottom μ-Dish 35 mm. Nuclei are in blue, OR-GFP protein is in green, vDNA is shown by intranuclear green spots of OR-GFP proteins.

Lastly, we attempted to visualize the viral DNA in live cells extracted from organs of HIV-1 infected animal models. Here we applied spinning disk technology to monitor viral DNA fluorescence spots in live CD4+ T cells freshly extracted from the bone marrow and spleen of BRGS/T A2DR2 ^OR-GFP^ HIS ANCH3-modified HIV-1 infected mice (Figure 6E). Dynamic 3D image acquisition occurred continuously (with a frame rate of 20 seconds), producing video recordings of infected cellsthat allowed us to easily distinguish infected CD4+ cells carrying potential intact viral DNA in the nucleus at 21 days post-infection (Figure 6E, movies 4, 5, 6, 7). Our results demonstrate that the HIV-ANCHOR^TM^ system represents a unique approach that enables real-time tracking of HIV-1 viral DNA directly within the nuclei of viable infected cells.

### Transcribing and silent HIV DNA detection in single cells during early stages of infection

Because ANCHOR^TM^ system is compatible with multiple imaging techniques(26, 32), we visualized HIV-1 transcripts emanating from ANCH3-modified HIV-1 genomes using single molecule inexpensive FISH (smiFISH). Twenty four unlabeled primary probes targeting the HIV-1 POL gene were prehybridized with fluorescently labeled secondary detector oligonucleotides via the shared FLAP sequence. Secondary probes were conjugated to Cy5. ANCHOR^TM^ technology paired with RNA smiFISH could discriminate between transcribing and non-transcribing HIV-1 DNA(32) and track down the transcribing proviruses of HIV-1 replicative strain in primary human CD4+ T cells (movie 8). CD4+ T cells were isolated from bone marrow, spleen, and lymph nodes of BRGST A2DR2^OR-GFP^ HIS mice after infection with ANCH3-modified HIV-1, and we observed viral DNA spots in 143 cells. The utilization of smiFISH enabled precise single-cell quantitative analysis of actively infected cells, identified as HIV RNA+ cells, in these organs (Figure 7A). As proviruses are considered highly transcribing forms with a vDNA-vRNA association or with vRNA signal in the cytoplasm in most acutely infected CD4+ T cells (25, 40), we further tracked transcriptionally active vDNA within the nucleus of these infected CD4+ T cells using 3D analysis (Figure 7A). A part of the analyzed CD4+ cells exhibited dual levels of vRNA expression, varying from vRNA present in both cytoplasm and nucleus, to specific instances of localized nuclear RNA foci, these foci were considered active proviruses. Some other vDNA signals that were not associated with vRNA signal in the nucleus and/or in the cytoplasm were considered low-transcribing proviruses or episomal forms characterized by minimal transcription activity (41). Notably, during the early stages of infection (21 days post-infection), some cells demonstrated no active transcription, despite containing vDNA (Figure 7A). A significant finding emerged when comparing maximal intensity value of the vRNA signal in the cytoplasm among CD4+T cells from different organs. We found that the vRNA max signal in the cytoplasm in the spleen cells was significantly higher than in cells derived from bone marrow or from lymph nodes (p value** <0.01; ****<0.0001) (Figure 7B). By imaging we set up an arbitrary threshold at a max vRNA value of 7500 (Figure 7B), since this signal did not give rise to a visible vRNA signal and we counted CD4+ T cells carrying an active vDNA and cells carrying a silent vDNA in the three analyzed organs (Figure 7C). Notably, we observed that the spleen exhibited an higher percentage of active vDNA (69%) in contrast to the bone marrow (56%) and lymph nodes (14%) (Figure 7B). Taken together the results in figures 7B and 7C imply that the spleen provides a favorable environment for extensive viral dissemination, setting it apart from the other two hematopoietic organs (bone marrow and lymph nodes) targeted by HIV. In the latter organs, the spread of the virus appears to be more constrained. Importantly, these data align with the greater viral spread observed in CD4⁺ T cells isolated from the spleen compared to those from the bone marrow of infected HIS mice (Figure 4C). This suggests an early establishment of cells with silent or low transcriptional activity of the cDNA, even during the acute phase, occurring more frequently in lymph nodes or bone marrow than in the spleen (Figures 7A,B,C).

**Figure 7.**
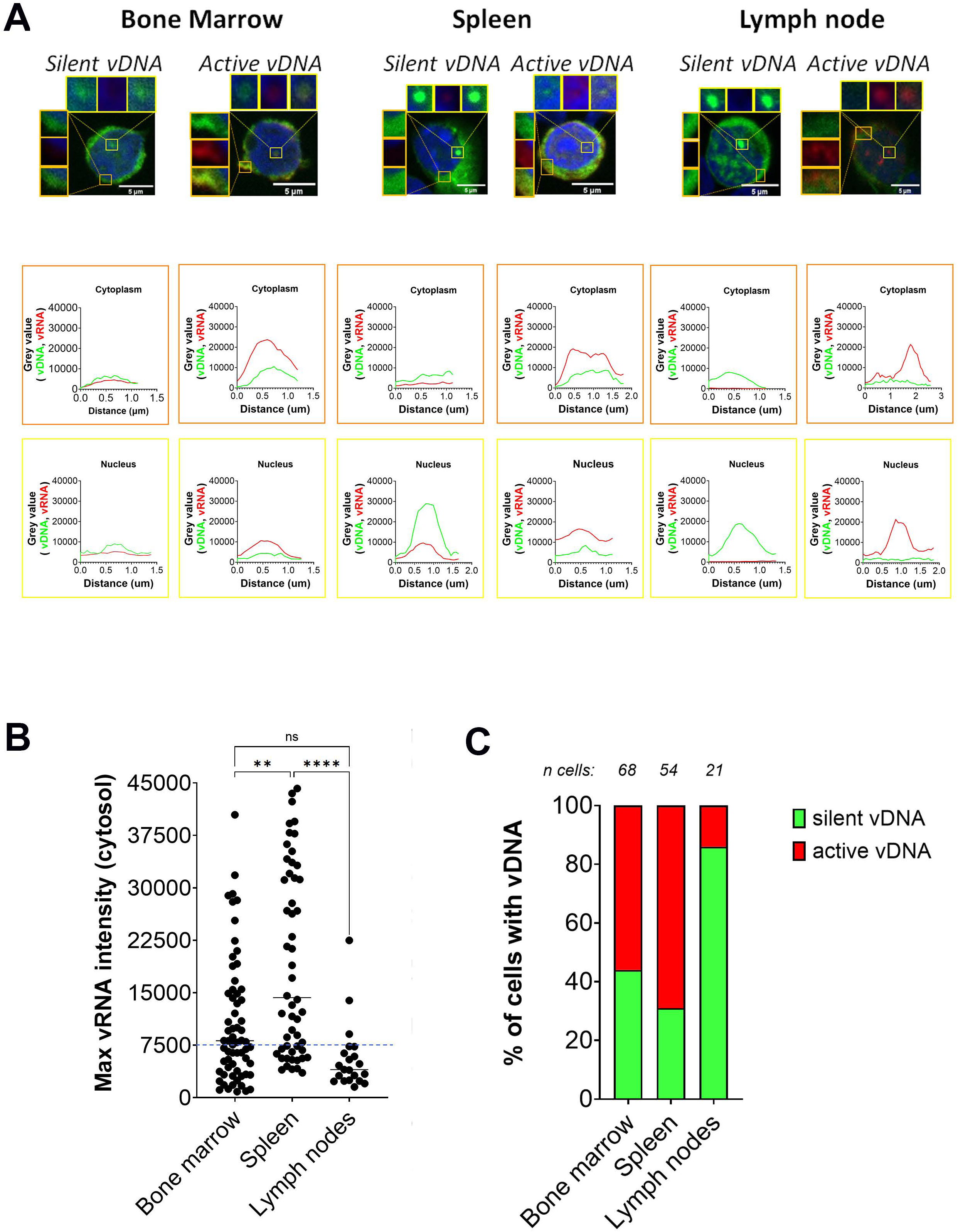
HIV-1 viral DNA detected in transcribing or silenced states in CD4⁺ T cells isolated from spleen, bone marrow, and lymph nodes of infected BRGS A2DR2 ^OR-GFP^ mice. **A,** Confocal imaging was performed with a 63X objective (Plan Apochromat, oil immersion, NA=1.4). We analyzed the 2D confocal images with the Fiji software version, and 3D confocal images were processed for multi-channel image splitting segmentation. Nuclei are labeled by Hoescht 33342 in blue, vDNA by OR-GFP in green, and vRNA by smiRNA-FISH in red. CD4+ T cells were sorted (d.p.i 21) from bone marrow, spleen and lymph nodes. A total of 2,616 CD4+ T cells from bone marrow, spleen, and lymph nodes were analyzed in 2D. vDNA spots were detected in 143 cells (5.47%). The distribution of transcriptionally active versus silent proviruses was estimated based on cytoplasmic vRNA signal intensity, with active vRNA defined as >7,500 arbitrary unit (A.U.) and silent as <7,500 A.U. **B,** Max vRNA intensities in the cytoplasm in single cells from bone marrow, spleen and lymph nodes are shown. Statistical test: ANOVA test, p values are shown in the graph. **C,** Percentage of cells carrying vDNA transcribing (>7,500 A.U.) and non-transcribing (<7,500 A.U.) in bone marrow, spleen and lymph nodes.

## Discussion

A better understanding of HIV-1 reservoir formation and persistance is crucial for the research on HIV-1 cure and an ability to identify these reservoirs *in vivo* holds immense potential for advancing this goal. One approach to achieve this involves making modifications to the HIV-1 genome to render it detectable through specific techniques. Establishment of this kind of system requires the alterations in the HIV-1 genome not to compromise the virus’s ability to replicate or its pathogenicity, as well as stability over successive replication cycles. Moreover, reliable detection of the modified virus in living cells without the need for cell fixation can open additional possibilities for understanding reservoir behaviour. Finally, this approach needs validation within a model system that accurately emulates the behavior of HIV and the formation of reservoirs in humans. In our study, we have introduced an innovative strategy that combines the ANCHOR^TM^ system with LV gene editing of hCD34+ cells in the BRGST HIS mouse model to address these three intricate experimental challenges.

In order to efficiently detect HIV-1 in infected cells, modifications to the viral genome must not impact on viral fitness nor disrupt the key stages of the HIV-1 life cycle: fusion with the cell’s membrane, reverse transcription, movement into the nucleus, integration of viral DNA into the host cell’s genetic material, and ultimately the production of infectious particles. This must be accomplished while ensuring a reliable detection method that avoids both false negatives and false positives. By employing the ANCHOR^TM^ system which does not rely on a direct fusion of protein with viral components, we mitigate the interference with the virus’s ability to infect the host cell. We observed that strains modified with ANCH3 effectively infected OR-GFP-transduced cell lines and primary activated CD4+ T cells *in vitro*. Furthermore, since the ANCH3 sequence can only be detected in its double-stranded DNA intranuclear form, it eliminates the possibility of signals from viral particles that cannot complete the aforementioned stages, unlike alternative methods like EdU and click chemistry (28). Additionally, when we compared the 400bp ANCH3 variant with the 1Kb version we observed that the ANCH3 1kb strain displayed reduced replication capabilities in primary CD4+ T cells *in vitro* and in the HIS mouse model *in vivo* and was not used in further experiments. This suggests that the extended ANCH3 configuration might have a detrimental effect on the packaging of viral RNA genomes and the assembly of viral particles, both of which are critical for viral fitness. Notably, we thoroughly characterized the 400 bp ANCH3 variant over the course of infection both in vitro and in vivo. We observed that a shorter form (250bp) remained stable throughout, and sequencing revealed that it retained four OR-GFP binding sites out of the original eight present in the ancestral construct. These four sites were sufficient to generate a bright, visible nuclear spot for visualizing viral dsDNA in both fixed and live cells. To corroborate these findings and assess the stability of this form over time, we generated a new viral construct carrying the naturally selected short ANCH3 sequence. Analysis by PCR and sequencing confirmed that its size remained unchanged throughout the course of infection. Altogether, these results indicate that the ANCH3 sequence does not interfere with reverse transcription, nuclear translocation, or viral packaging. Importantly, we observed that the replication efficiency of the tagged virus was comparable to that of the wild-type virus. CD4⁺ T cells isolated from both the spleen and bone marrow were capable of producing and spreading the virus under cell culture conditions. Intriguingly, we found that spleen-derived cells exhibited a greater potential for viral spread compared to those derived from the bone marrow. In the early stages of acute infection, CD4+ T cells carrying the HIV-1 genome have the potential to disseminate to various organs and undergo transcriptional silencing, transforming into latent, long-lived reservoirs with the capacity for later reactivation (5). Consequently, any detection system designed for the investigation of the reservoirs must undergo testing in a model capable of mimicking some aspects of reservoir formation. The utilization of BRGS/T A2DR2 mice xenografted with hCD34+ cells, genetically modified with a LV carrying OR-GFP, proved highly effective in achieving robust engraftment, reconstitution, and development of key immune populations, notably CD4+ T cells in crucial anatomical sites such as the bone marrow, spleen, and lymph nodes. This reconstitution maintained the expression of OR-GFP under the EF1a promoter, ensuring the fidelity of our model’s representation. Our approach allowed for the differentiation of individual viral DNA molecules within CD4+ T cells obtained from various organs of the HIS mice during early stages of infection. A combination with RNA smiFISH technique enabled the detection of both transcribing and non-transcribing viral DNA molecules. We considered cells with both vDNA foci and cytoplasmic vRNA signals as actively transcribing, whereas cells showing only vDNA foci were classified as carrying silent forms, potentially representing newly formed reservoirs with suppressed transcription. Additionally, we have observed that CD4+ T cells derived from the bone marrow and lymph nodes display a higher tendency to harbor the silent viral DNA form within their nuclei. While HIV-1 shows a strong preference for integrating into euchromatin, often within introns of highly expressed genes, during acute infection, comprehensive cross-sectional and longitudinal studies have revealed that intact proviruses accumulate in chromosomal regions characterized by heterochromatin feature (42). Alternatively, both observations may stem from the episomal viral DNA, a substantial component of viral DNA during acute infection. Although the episomal viral DNA form is generally considered non-transcribing, there are indications suggesting that it can undergo transient transcription, potentially supporting latency, virus production, and immune responses (43–45), offering a plausible explanation for both silent and transcribing detected viral DNA. Currently, our existing data does not definitively distinguish between these forms when ANCHOR^TM^ alone is utilized. Animal models offer unique advantages over *in vitro* systems and human samples, allowing for a more comprehensive investigation of HIV-1 and enabling preclinical evaluation of drugs and vaccines. Among these models, HIS mice provide significant benefits over non-human primates, including higher reproductive rates, the ability to be infected with unmodified wild-type HIV-1, cost-effectiveness in maintenance, and the use of genetically identical strains (38, 46). However, HIS mice differ from humans in aspects of immune cell development and repertoire. Advances in transgenic mouse technology have helped mitigate some of these differences. For instance, the induction of TSLP expression in the BRGS model restores secondary lymphoid tissues, enhances T follicular helper cell development, and improves the humoral immune response, thereby offering a more accurate representation of the human immune system (35). Despite these improvements, HIS mice still exhibit a higher CD4:CD8 ratio and distinct CD4+ T cell subset distribution (naïve, central memory, and effector memory) compared to humans (47). Combined with their underdeveloped mucosal immune system, these factors can alter the course of CCR5-tropic (R5) HIV-1 infection, leading to a latency landscape different from that in humans (47, 48). Additionally, while HIV-1 reservoirs do form in the brains of HIS mice (38, 49) differences in the presence of microglial cells—key targets for HIV-1 in the human brain—may limit the usefulness of HIS models for studying HIV pathology in the brain (50). Despite these limitations, HIS mice have been instrumental in advancing our understanding of HIV transmission, pathogenesis, prevention, treatment, and potential cures, with many findings later confirmed in humans and/or non-human primates. The integration of the ANCHOR^TM^ system with BRGST mice has enabled the detection of HIV-1 DNA in CD4+ T cells from various organs, employing techniques such as confocal microscopy and live imaging using a spinning disk. These technologies offer a platform for in-depth exploration of viral reservoirs, especially under combined antiretroviral therapy (cART), facilitating analysis of the tissue-specific and cell type-specific distribution of viral DNA and viral RNA and shedding light on the dynamics of HIV persistence in different physiological niches.

Moreover, the BRGST^OR-GFP^-ANCHOR^TM^ system provides an opportunity to investigate the effects of latency reversal agents (51), the therapies aimed at targeting and reactivating latent virus reservoirs, leading to their exposure to the immune response or other treatments. Understanding how these agents influence the reservoir landscape within distinct tissues and cell types will be instrumental in refining therapeutic strategies. Additionally, the role of host factors in HIV pathobiology remains a focal point, and can be addressed with the application of single-cell RNA sequencing (scRNA-seq) on isolated viral DNA-containing cells, offering an approach to investigate the complexity of host-virus interactions at the molecular level. On the other hand, our approach can be instrumental in gaining a deeper understanding of the mechanism of action of the new promising drugs that target the initial stages of the viral life cycle as well as FDA approved drugs such as Lenacapavir(52, 53). Furthermore, our HIS-ANCHOR™ system could be used for future studies monitoring viral rebound without the need for in vitro passaging, which could otherwise alter the physiological process of viral reactivation. n the context of *in vivo* experimentation, the intranuclear localization of viral DNA holds the potential to significantly enrich our understanding of cellular host factors and the intricate organization of chromatin. A strategic approach involving the labeling of viral DNA, ideally in living cells or in fixed specimens, stands to furnish a more comprehensive dataset concerning kinetics, precise localization, and interactions within the nucleus. Moreover, this approach is poised to illuminate the trajectory of infected cells, contingent upon the distinct phases of HIV infection.

## Materials and methods

### Cell lines

All media were supplemented with 10% Fetal Bovine Serum (FBS) and 100 U/mL penicillin-streptomycin (Pen/Strep). We kept all cells in an incubator at 37°C and 5% CO2. HEK 293T cells (ATCC) are human embryonic kidney cells that produce lentiviral vectors and HIV-1. HeLa P4R5 are cervical carcinoma cells carrying the β-galactosidase gene under HIV-1 LTR promoter and expressing CD4, CXCR4, and CCR5. HEK293T and HeLa P4R5 cells were cultivated in Gibco Dulbecco’s Modified Eagle Medium (DMEM).

### Human primary cells

Peripheral Blood Mononuclear Cells (PBMCs) were isolated through density gradient centrifugation with Ficoll 400 from healthy donors’ blood, provided by EFS (Etablissement Français du Sang, Paris). Primary CD4+ T cells were freshly positively selected with human CD4 Microbeads (Miltenyi Biotec #130-045-101) from PBMC. Fetal liver CD34+ cells (Advanced Bioscience Resources Inc., USA) were isolated with CD34+ affinity matrices according to the manufacturer’s instructions (Miltenyi Biotec). Cord blood CD34+ cells were isolated from Cord-blood (Cell therapy laboratory in Saint Louis Hospital, APHP, Paris) through density gradient centrifugation, purification, and enrichment by positive selection using indirect human CD34 MicroBead Kit, (Miltenyi Biotec #130-046-701)(54) Isolated hCD34+ cells were stored in in vapor-phase liquid nitrogen below –135°C.

### Lentiviral Vectors

OR-GFP cDNA was cloned under control of the long (1239 bp) or short (250 bp) EF1a promoters inserted between MluI and BamHI sites. The pFLAPOR-GFP(26, 55) plasmid contained also a mutated WPRE (Woodchuck Posttranscriptional Regulatory Element) sequence to improve protein expression. ANCH3 sequence (400bp or 1Kb) was cloned in the sites MluI and XmaI of NL4-3 IRES-eGFP Infectious Molecular Clone (pBR43IeG-NA7nef) (NIH) at the place of iresGFP. HIV AD8 ANCH3sh (250 bp) construct was obtained performing a digestion of the HIV AD8 ANCH3 (400 bp) plasmid with the restriction enzyme XmaI. After the run on agarose gel, the extraction and purification, the construct was ligated with T4 Ligase (Thermo Scientific) and amplified in competent DHα E. coli. Both bacterial strains were routinely grown in Luria-Bertani (LB) medium at 37^°^C (or at the indicated temperatures) while shaking at 200 RPM. Plasmids were quantified with a NanoDrop 2000c spectrophotometer (Thermo Fisher Scientific), aliquoted, and stored at −20°C. Plasmid DNA was verified by enzymatic digestion and sequencing of the region proximal to the transgene insertion sites or the full plasmid has been sequenced. We produced Non-replicative LV in Human Embryonic Kidney (HEK)-293T cells. Lentiviral particles were produced by transient calcium phosphate co-transfection of HEK293T cells with the pflap vector plasmid (pFlap EF1a (long or short) OR(3)-GFP), a vesicular stomatitis virus G (VSV-G) Indiana envelope plasmid and an encapsidation plasmid. Supernatants were harvested 48h post-transfection, clarified, concentrated by ultracentrifugation, aliquoted, and stored at −80°C. Vector titers were determined by transducing 293T cells. The titer, proportional to the efficacy of nuclear gene transfer, is defined as Transduction Unit (TU)/mL by qPCR on total lysates at day 3 post-transduction, by use of forward 5’-TGG AGG AGG AGA TAT GAG GG-3’ and reverse 5’-CTG CTG CAC TATACC AGA CA-3’ primers, specific to pFLAP plasmid and forward and, reverse 5’-CTG CTG CAC TATACC AGA CA-3’ primers specific to the host housekeeping gene CD3 previously described.

### Transduction of Human CD34+ cells

After thawing, hCD34+ cells were pre-activated overnight and transduced. The transduction medium was Xvivo 20 (Lonza, Basel, Switzerland) supplemented with SCF 300 ng/mL (peprotech), Flt3-L 300 ng/mL (peprotech), TPO 100 ng/mL (peprotech) and IL3 10-60 ng/mL(peprotech). We used non-toxic enhancers of transduction 4 μg/mL Protamine sulfate (Sanofi-Aventis) at a concentration of 4 ug/mL and PGE2 (Sigma) at a concentration of 10ug/ml. Culture plates were coated with fibronectin CH-296 (RetroNectin; Takara, Shiga, Japan) for the initial experiment.

### NL 4.3 AD8 replication competent HIV tag

We produced the replicative HIV-1 constructions in Human Embryonic Kidney (HEK)-293T cells. The titer, proportional to the efficacy of HIV DNA integration, was determined as Transduction Unit (TU)/mL by qPCR on total lysates at 48h post-infection in HelaP4R5, by use of forward 5’-TGG AGG AGG AGA TAT GAG GG-3’ and reverse 5’-CTG CTG CAC TATACC AGA CA-3’ primers, specific to U5 LTR.

### HIV infection assay in Hela P4R5

For imaging studies in adherent cell lines, the cells were plated on coverslips (12mm diameter in 24 wells plates, ThermoFisher) at least 24h before fixation. The Hela P4R5 cell line was plated on D1 at 10^5 cells on DMEM medium supplemented with 10% FCS and 2 mmol/L L-glutamine on a coverslip from 24 wells plate. Hela P4R5 was transduced with a multiplicity of infection (MOI) 1 of LV EF1aOR-GFP on day 2. After 24 hours of transduction, Hela P4R5 were infected with MOI 50 of NL4.3 AD8 ANCH 1Kb and ANCH 400bp for 30 hours. All cells were washed with PBS and fixed with 4% PFA for 15 minutes 5. Nuclear OR-GFP dots from integrated and episomal forms were counted by confocal after Hoechst 33342 epifluorescence staining 1:10000 (Invitrogen #H3570). Coverslips were mounted on glass slides (Star Frost) with Prolong Diamond Antifade Mountant (Life Technologies #P36970).

### OR-GFP transduced CD4+ T cell HIV infected assay

Primary human CD4+ T cells were isolated using positive magnetic sorting (CD4 MicroBeads, human 130-045-101 Miltenyi-Biotec). Human CD4+ T cells were activated for 72 hours, using Activation/Expansion Kit (Miltenyi Biotec, #130-091-441). Activated primary human CD4+ T cells were transduced with OR-GFP lentiviral vector (MOI 10) and infected with MOI 20 of ANCH3 tagged virus. After 3 days of infection, suspension CD4+ T cells were plated on polylysine, washed with PBS, and fixed with 4% PFA for 15 minutes. Nuclear OR-GFP dots from integrated and episomal viral forms were counted by confocal after Hoechst 33342 epifluorescence staining 1:10000 (Invitrogen #H3570). Coverslips were mounted on glass slides (Star Frost) with Prolong Diamond Antifade Mountant (Life Technologies #P36970).

### p24 assay

CD4+ T cells were purified from PBMC and were isolated using positive magnetic sorting (CD4 MicroBeads, human 130-045-101 Miltenyi-Biotec). The day after sorting, CD4+ T cells were activated with T Cell Activation/Expansion Kit, human (130-091-441, Miltenyi-Biotec) and human IL-2 (400 IU/ml Peprotech, #17881063in RPMI-1640 medium supplemented with GlutaMAX-I (Gibco), 10 % serum, 1% penicillin-streptomycin (v/v). At 3 days of activation, CD4+ T cells were infected with HIV NL4.3 AD8 WT, HIV NL4.3 AD8 ANCH 400bp or HIV NL4.3 AD8 ANCH 1000bp (MOI 0.1) in the 96 wells plate at the concentration of 0,2×106 cells/200ul in RPMI-1640 medium supplemented with GlutaMAX-I (Gibco), 10 % serum, 1% penicillin-streptomycin (v/v), human IL-2. The supernatants were collected at days 3; 7; 10; 14; 21 and after the collect of supernatant, we change the medium with the new RPMI-1640 medium supplemented with GlutaMAX-I (Gibco), 10 % serum, 1% penicillin-streptomycin (v/v), human IL-2. Secreted P24 levels were measured from culture supernatants diluted 1/1000 with Lenti-X p24 Rapid Titer kit (632200 Takara) following manufacturer’s instructions.

### RT qPCR 5’LRT and ANCH (400bp/ 1KB) in plasma

Total viral RNA was extracted from the plasma of infected mice via the QIAamp Viral RNA Mini Kit (Qiagen, Hilden Germany) following the manufacturer’s protocol. Total viral RNA was eluted into 60 μL of DEPC. RNA copies for each sample were determined using the qRT-PCR SuperScript III Platinium (ThermoFisher, #11732-020), and absolute RNA copies were calculated from a standard curve run in parallel using lentiviral ANCH 400 bp or 1Kb genomic RNA quantified at 40 ng/ uL. Real-Time RT-PCR (qRT-PCR) reactions were duplicated on specific genomic regions: LRT, ANCH400bp, and ANCH 1Kb.

### Reverse transcription and PCR ANCH3 in CD4+ cells supernatant

Activated CD4+ T cells derived from two different donors were infected with NL4.3 HIV-1 AD8 ANCH3 (400 bp) (MOI 0.5). After 21 days p.i., the supernatant was collected and the total viral RNA was extracted using QIAmp Viral RNA mini Kit (Qiagen, Hilden Germany) and following the manufacturer’s protocol. 40 ng of RNA copies were used to perform the reverse transcription with Maxima H Minus Reverse Transcriptase (Thermoscientific) protocol and reagents, following the manufactures’ protocol. Samples were incubated for 30 minutes at 50°C and then exposed to 85°C to deactivate the enzyme. Afterwards, the obtained cDNA was amplified using PCR cycles (98°C for 10 minutes, 30 cycles of 98°C for 1 sec, 56°C for 5 sec, 72°C for 10 sec, at the end 10 minutes at 72°C and storage). The amplified cDNA was then run on a 1.3% agarose gel and visualized through UV light exposure.

### Amplification of ANCH3 sequence in Primary CD4+ T cells supernatant and in mice plasma

The reverse transcription was performed using Maxima H Minus Reverse Transcriptase (EP0752, Thermo Scientific) protocol and reagents. 100 ng of RNA were reverse transcribed using primers that anneal on the viral genome immediately before and after the site of cloning of the ANCH3 cDNA: 5’ AGGAAGAACAAGACCCAGTAAAACG (Forward) and 5’ GTGAATTAGCCCTTCCAGTCCC (Reverse) for the samples from Primary CD4+ T cells supernatant. A first round PCR using primers annealing on the viral genome were used for the samples from mice plasma 5’ GACTGCATCCAGAGTTCTTT (Forward) and 5’ GTGAATTAGCCCTTCCAGTCCC (Reverse). 5ul of cDNA were amplified with the same primers used for the reverse transcription and following the protocol of Phusion Flash High-Fidelity PCR Master Mix (F-548S, Thermo Scientific). The PCR steps were performed as follows: 98°C for 10 minutes, 30 cycles of 98°C for 1 sec, 60°C for 5 sec, 72°C for 10 sec, at the end 10 minutes at 72°C and storage at 4°C. For the PCR nested, required in samples from mice plasma, the primers used were: 5’ AGGAAGAACAAGACCCAGTAAAACG (Forward) and 5’ TCTTTTAAAAAGTGGCTACC (Reverse), with same PCR steps as previously explained. The amplified cDNA was then run on agarose gel and visualized through UV light exposure.

### HIV RNA Labelling

To visualize individual HIV vRNA molecules, we used the smiFISH approach. Twenty four unlabeled primary probes were designed to target the HIV-1 POL gene. They could be prehybridized in vitro with fluorescently labeled secondary detector oligonucleotides via the shared FLAP sequence. Secondary probes are conjugated to Cy5. Cells were fixed, washed twice with PBS, and stored in nuclease-free 70% ethanol at −20°C until labeling. On the day of the labeling, the samples were brought to room temperature, washed twice with wash buffer A (2× SSC in nuclease-free water) for 5 min, followed by two washing steps with washing buffer B (2× SSC and 10% formamide in nuclease-free water) for 5 min. The reaction mixture contained primary probes at a final concentration of 40 pm and secondary probes at a final concentration of 50 pm in 1× NEB3 buffer. Pre-hybridization was performed in a PCR machine with the following cycles: 85°C for 3 min, followed by heating to 65°C for 3 min, and a further 5 min heating at 25°C. 2 µl of this FISH-probe stock solution was added to 100 µl of hybridization buffer (10% (*w*/*v*) dextran, 10% formamide, 2× SSC in nuclease-free water). Samples were placed on Parafilm in a humidified chamber on 100 μl of hybridization solution, sealed with Parafilm, and incubated overnight at 37°C. The next day, cells were washed in the dark at 37°C without shaking for > 30 min twice with pre-warmed washing buffer B. Samples were washed once with PBS for 5 min, stained with Hoescht 33342 in PBS (1:10,000) for 5 min, and washed again in PBS for 5 min. Samples were mounted in ProLong Gold antifade mounting medium. Probes for RNA FISH are described in (29, 32).

### Animal care

All experiments involving live animals were approved by an ethical committee at the Institut Pasteur (CETEA-2013-0131) and validated by the French Ministry of Education and Research (Reference # 02162.01). *In vivo* studies were conducted at the Institut Pasteur Animalery (A3). Animals were housed in isolators under pathogen-free conditions with humane care and were anesthetized with isoflurane to minimize suffering.

### HIS mice model

For the ANCHOR^TM^ *in vivo* fitness validation, BRGS/T A2DR2 HIS mice were generated as previously described [29, 34]. Briefly, human fetal liver (Advanced Bioscience Resources Inc.) CD34+ cells were isolated using MicroBead Kits (Miltenyi). CD34, CD38, and HLA-A2 expressions were phenotyped by FACS and the HLA class II allele haplotype analysis by PCR (LABType SSO, One Lambda). 7–15 × 10 [35] CD34+CD38− cells from HLA-A*02+ HLA-DRB1*15+ donors, or from 1:1 mixtures of HLA-A*02+ and HLA-DRB1*15+ donors, were intrahepatically injected into 2–10-day-old mice that had been sublethally irradiated (3 Gy). For other experiments, irradiated pups were injected with CD34+ isolated cell human cord-blood. The efficiency of humanization in the blood was assessed by flow cytometry using the following formula: %hCD45 × 100/(%hCD45 + %mCD45); Once the engraftment of mice was confirmed, they were challenged with either 10^5^ TCID50 of WT NL4.3AD8 or 10^5^ TCID50 NL4.3 AD8HIV-1 ANCH3 strains. Every week, blood was collected from each animal, and the HIV plasma viral load was determined by qRT-PCR. Animal weight was monitored before HIV infection and then once weekly until sacrifice.

### Mice tissue collection

To obtain single cell suspensions LNs (all visible LNs of BRGST A2DR2^OR-GFP^ mice) were pooled, minced and filtered through a 100-μ m cell strainer (Falcon). Spleen was processed in the same manner. Femur and tibia bones were crushed with a mortar and pestle and filtered through a 100-μ m cell strainer (Falcon). Erythrocytes in spleen and bone marrow were lysed with Hybri-Max red blood cell lysing buffer (Sigma, R7757). All cell-preparation steps were performed using RPMI 1640 Glutamax (Life Technologies), 100 U/mL penicillin, and 100 g/mL streptomycin (Invitrogen). All cell suspensions were counted with Trypan blue (Life Technologies). CD4+ cells were freshly positively selected with human CD4 Microbeads (Miltenyi Biotec #130-045-101) from splenocytes, bone marrow, and lymph nodes. Suspension of CD4+ cells were plated on polylysine, washed with PBS, and fixed with 4% PFA for 15 minutes.

### Infection of HeLa P4R5 cells using the supernatant isolated from CD4+ cells from spleen and bone marrow of infected mice

CD4⁺ T cells from BRGS mice infected with HIV/NL4.3 AD8 ANCH 400pb were freshly isolated from splenocytes and bone marrow using human CD4 MicroBeads (Miltenyi Biotec, #130-045-101), as previously described in the “Mice tissue collection” section of the Materials and Methods. Cells were cultured in RPMI-1640 medium supplemented with GlutaMAX-I (Gibco), 10% fetal calf serum, and 1% penicillin-streptomycin (v/v).

The day after sorting, CD4⁺ T cells were activated using the Human T Cell Activation/Expansion Kit (Miltenyi Biotec, #130-091-441) and human IL-2 (100 IU/mL; Peprotech, #17881063) at a density of 2 × 10⁶ cells/mL in the same supplemented RPMI-1640 medium, and incubated for three days.

5 × 10³ HeLa P4R5 cells were seeded in 96-well flat-bottom plates (ThermoFisher) in DMEM supplemented with 10% FCS and 2 mM L-glutamine. The following day, cells were infected with 100 µL of filtered (0.44 µm) supernatant collected from the activated CD4⁺ T cells. After 24 hours, the supernatant was removed, and HeLa P4R5 cells were washed twice with PBS and cultured in fresh DMEM medium (supplemented as above) for an additional 48 hours.

On day 3, supernatants were collected to measure the total viral RNA by RT-qPCR. On the same day, infectivity of HeLa P4R5 cells was assessed by quantifying β-galactosidase activity using a chemiluminescent β-Gal Reporter Gene Assay (Roche, #11758241001), and normalized to total protein content.

### Flow Cytometry

Whole blood collected in EDTA-coated Microvettes (Sarstedt) was directly stained with fluorochrome-conjugated antibodies (Table S1) at 4°C for 30 min, erythrocytes were lysed, and samples fixed with FACS Lysing/Fixation solution (BD #349202). For the organ-derived cells, a maximum of 2 × 10^6^ cells were used for analysis, prior to staining they were incubated for 10 min at RT with IgG from human serum (Sigma-Aldrich, #I2511) and FcR blocking reagent Mouse (Miltenyi, #130-092-575). For surface marker staining, respective fluorochrome-conjugated antibodies (Table S1), for dead cell exclusion Viability dye eFluor 506 (Invitrogen) and cell number determination CountBright absolute counting beads (Invitrogen), were diluted in brilliant stain buffer (BD). Staining was done at 4°C for 30 min, after washing cells were fixed in 4% PFA (EMS) and resuspended for acquisition. Washing steps were done in FACS buffer (1× DPBS (Gibco) with 2% fetal calf serum (eurobio), 2 mM EDTA (Invitrogen)). For intra cellular staining cells were fixed by Cytofix/Cytoperm (BD) and subsequently incubated with intra cellular antibody diluted in Perm/Wash (BD) for 30 min at 4°C. Data were acquired on an LSR Fortessa (BD) or Attunes and analyzed with FlowJo 10 software (FlowJo LLC).

### Confocal microscopy

Confocal microscopy was carried out with a Zeiss inverted LSM700 microscope, with a 63X objective (Plan Apochromat, oil immersion, NA=1.4). We analyzed the 2D confocal images with the Fijii software version, and 3D confocal images were processed for multi-channel image splitting, Hoescht 33342, vDNA, and vRNA segmentation.

### Live imaging Time lapse microscopy

Primary CD4+ T lymphocytes were plated in a polymer-bottom dish (ibidi #81156) coated with collagen IV (sigma #C6745), as manufacturer protocol, for aquisition 2 h later. Specifically, CD4+ T lymphocytes from the spleen of infected mice were imaged for about 6 minutes in continuous in 3D (stack spacing 0.2 µm) with an UltraView VOX inverted microscope (PerkinElmer), based on a CSU-X spinning-disk (Yokogawa), and using a 63X objective (Plan Apochromat, oil immersion, NA=1.4).

### Statistical Analysis

Data are presented as mean and s.e.m. Sample size and the statistical tests used for each experiment are described in the figure legends. No statistical methods were used to pre-determine sample size. Statistical analysis was performed using GraphPad Prism software.

## Ethics statement

Animals were maintained in isolators under pathogen-free conditions and received humane care throughout the study. To minimize discomfort, anesthesia was performed using isoflurane. All procedures related to the generation and characterization of humanized mice complied with established ethical guidelines. The experimental protocol was reviewed and approved by the Institutional Animal Care and Use Committee at the Institut Pasteur (CETEA-2013-0131) and subsequently authorized by the French Ministry of Education and Research (Reference #02162.01).

## Acknowledgments

The following reagent was obtained through the NIH HIV Reagent Program, Division of AIDS, NIAID, NIH: Human Immunodeficiency Virus Type 1 NL4-3 IRES-eGFP Infectious Molecular Clone (pBR43IeG-NA7nef), ARP-11352, contributed by Drs. Jan Münch, Michael Schindler and Frank Kirchhoff. We are grateful to ANRS, FRM and Sidaction grants and Institut Pasteur in Paris.

