## Supplementary file for "Tracking HIV-1 DNA fate from Cell Culture to Humanized mice Tissues"

| BRGST OR-GFP | TSLP expression | CB donors | Purity CD34+ (2 passages) | CD34+ CD38-% | Transduction MOI | CD34+ injected/ Mouse | Age (days) of infusion (pups) | ALU integration/ cell | %CD34+ ORGFP+ (D7) |
|---|---|---|---|---|---|---|---|---|---|
| 1 | BRGS | CBa4 | 96.6 | 3.60% | 50 | 2.00E+05 | 7 | 1.4 | 66.8 |
| 2 | BRGS | CBa4 | 96.6 | 3.60% | 50 | 2.00E+05 | 7 | 1.4 | 66.8 |
| 3 | BRGS | CBa4 | 96.6 | 3.60% | 50 | 2.00E+05 | 7 | 1.4 | 66.8 |
| 4 | BRGS | CBa4 | 96.6 | 3.60% | 50 | 4.00E+05 | 7 | 1.4 | 66.8 |
| 5 | BRGS | CBa4 | 96.6 | 3.60% | 50 | 4.00E+05 | 7 | 1.4 | 66.8 |
| 6 | BRGS | CBa4 | 96.6 | 3.60% | 50 | 4.00E+05 | 7 | 1.4 | 66.8 |
| 7 | BRGS | CBa4 | 96.6 | 3.60% | 50 | 4.00E+05 | 7 | 1.4 | 66.8 |
| 8 | BRGS | CBa7 | 99.1 | 35.1 | 50 | 4.00E+05 | 4 | 0.8 | 77.2 |
| 9 | BRGS | CBa7 | 99.1 | 35.1 | 50 | 4.00E+05 | 4 | 0.8 | 77.2 |
| 10 | BRGS | CBa7 | 99.1 | 35.1 | 50 | 4.00E+05 | 4 | 0.8 | 77.2 |
| 11 | BRGST | CBa7 | 99.1 | 35.1 | 50 | 4.00E+05 | 4 | 0.8 | 77.2 |
| 12 | BRGST | CBa7 | 99.1 | 35.1 | 50 | 4.00E+05 | 4 | 0.8 | 77.2 |
| 13 | BRGST | CBa5 | 93.2 | 80.9 | 50 | 4.00E+05 | 4 | - | 69.3 |
| 14 | BRGST | CBa5 | 93.2 | 80.9 | 50 | 4.00E+05 | 4 | - | 69.3 |
| 15 | BRGST | CBa6 | 99.1 | 50.9 | 50 | 4.00E+05 | 4 | 0.4 | 60.1 |
| 16 | BRGST | CBa6 | 99.1 | 50.9 | 50 | 4.00E+05 | 4 | 0.4 | 60.1 |
| 17 | BRGST | CBa6 | 99.1 | 50.9 | 50 | 4.00E+05 | 4 | 0.4 | 60.1 |
| MEDIAN | | | 96.6 | 35.1 | 50 | 4.00E+05 | 4 | 0.8 | 66.8 |
| Min | | | 93.2 | 0.036 | 50 | 2.00E+05 | 4 | 0.4 | 60.1 |
| Max | | | 99.1 | 80.9 | 50 | 4.00E+05 | 7 | 1.4 | 77.2 |

**Table S1. Recapitulative data on mice xenotransplanted.** CB donors, purity of CD34+, MOI of transduction, number of CD34+ cells injected per mouse, number of proviruses (ALU PCR), % of OR-GFP CD34+ positive cells injected 24h post transduction and data at day 7 post transduction transduced cells kept in culture.

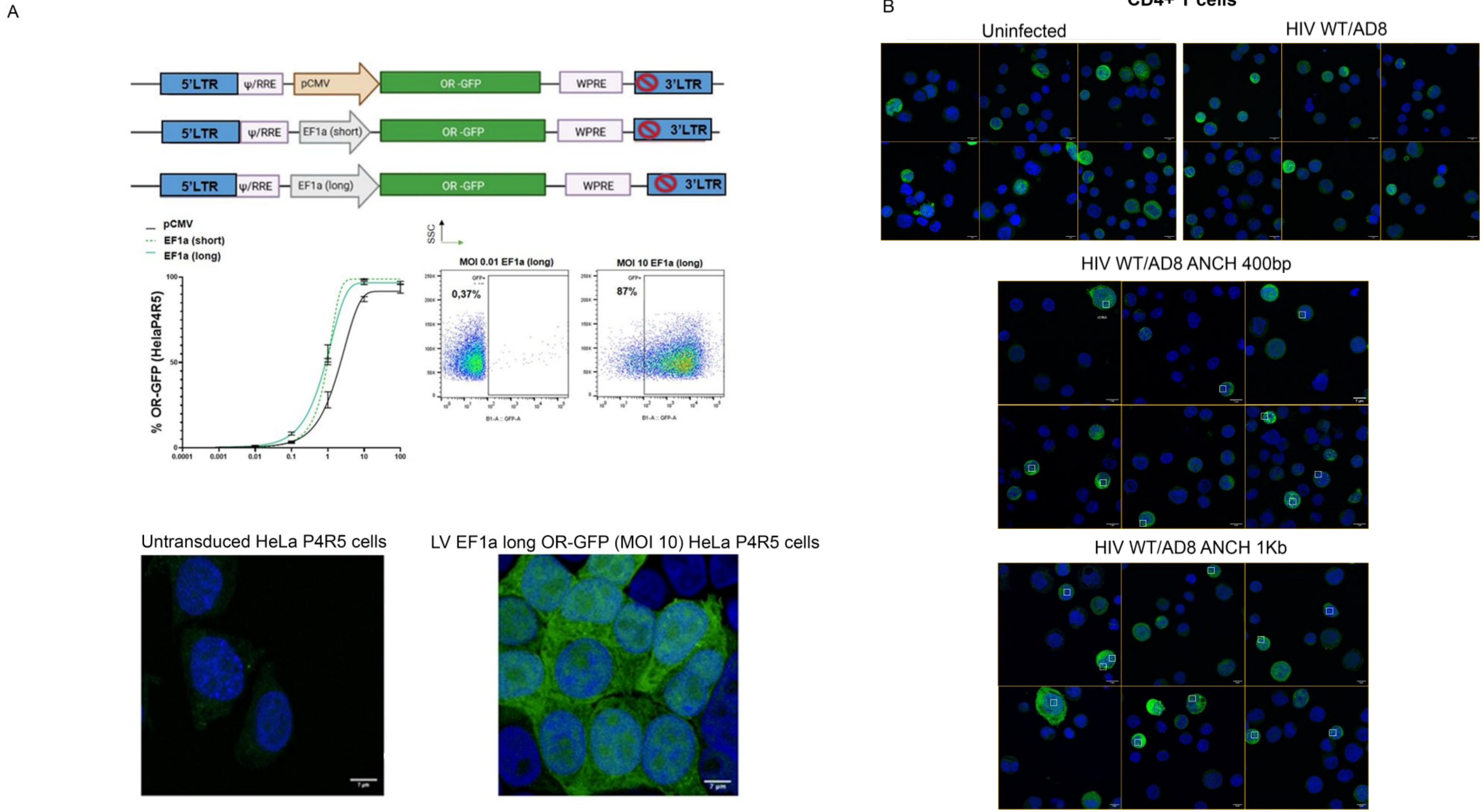


**Fig. S1. HIV-1 ANCHOR system to detect ds vDNA in the nuclei of target cells.** (A) On the top, designs of constructs of LVs used. Percentage of OR-GFP positive HeLa P4R5 cells transduced at different MOIs (x axes) with the three LVs. On the right example of FACS analysis. On the bottom example of HeLa P4R5 transduced or not by confocal (OR-GFP in green and nuclei in blue). **(B)** Primary CD4+T cells expressing OR-GFP uninfected or infected with HIV WT AD8 or with HIV WT/AD8 ANCH 400bp or with HIV WT/AD8 ANCH 1Kb. OR-GFP is mainly expressed in the cytoplasm but the bright spots in the nucleus identify vDNA, especially at the highest MOI, nuclei are labelled by Hoechst in blue. Scale bar 7µm.

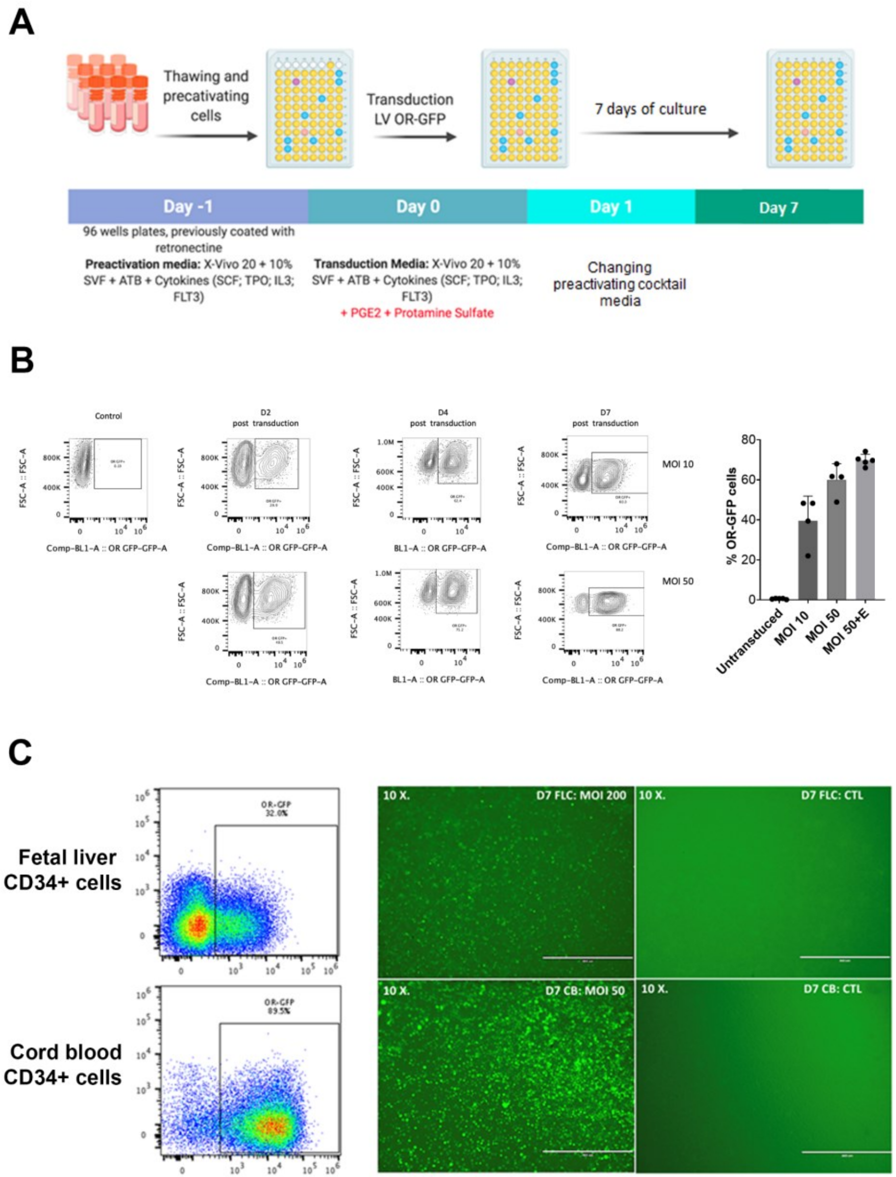


D

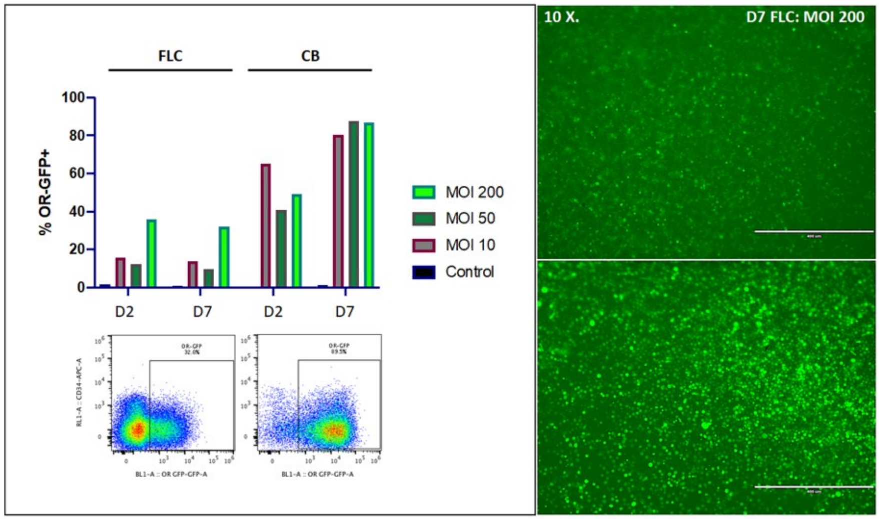


**Fig. S2. Optimization protocol for an efficient transduction of hCD34+ cells with LVs EF1along OR-GFP. (A)** Optimized transduction protocol for hCD34+ cells derived from cord blood. **(B)** FACS analysis and results showed in the bar graph on the right of hCD34+ cells transduced with different MOIs. **(C)** comparison of efficiency of transduction between CD34+ cells derived from fetal liver (top) and cord blood (bottom) using the protocol described in A. **(D)** Cytofluorimetry analysis (left) and fluorescence microscopy to compare the transduction of fetal liver-derived CD34+ cells and cord blood-derived CD34+ cells using different MOIs of LV and different time post-transduction.

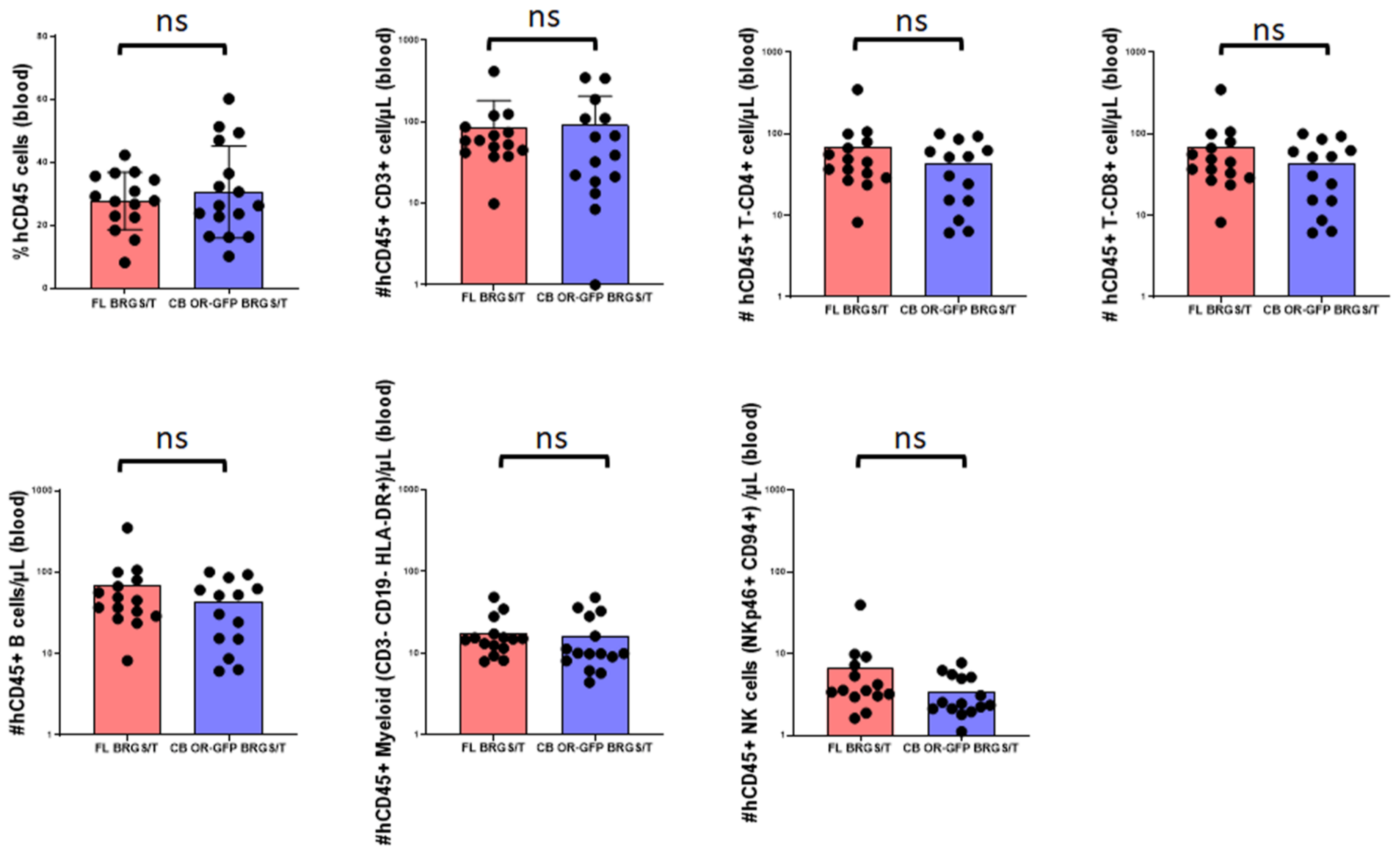


**Fig. S3. Humanization efficiency.** Calculated via cytofluorimetry comparing mice xenografted with fetal liver-derived (FL) CD34+ cells and cord blood-derived (CB) CD34+ cells.

A

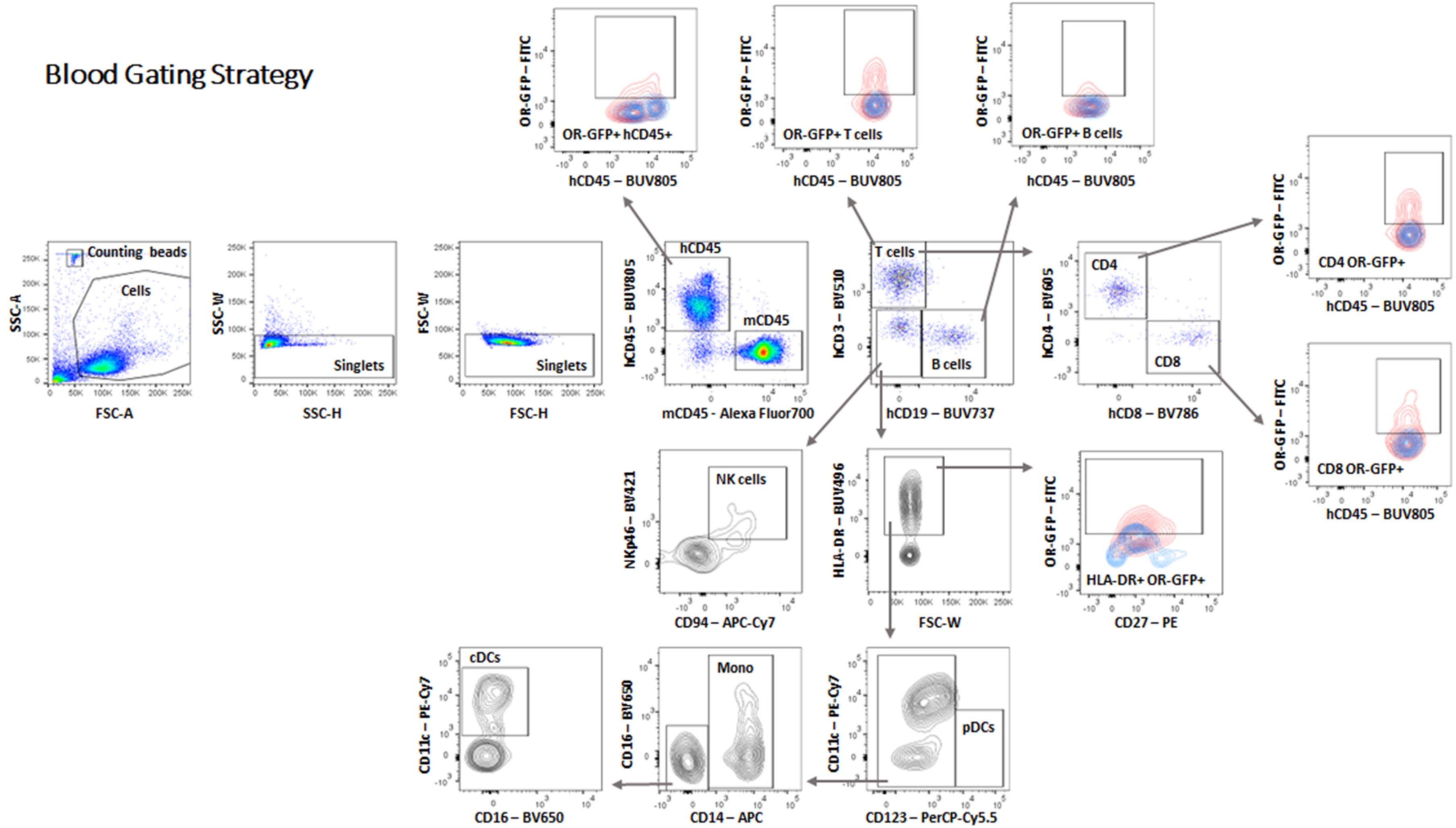


**Fig. S4. Gating strategy for FACS analysis.** Immune cell subsets and representative FACS plots for OR-GFP signal in blood.

B

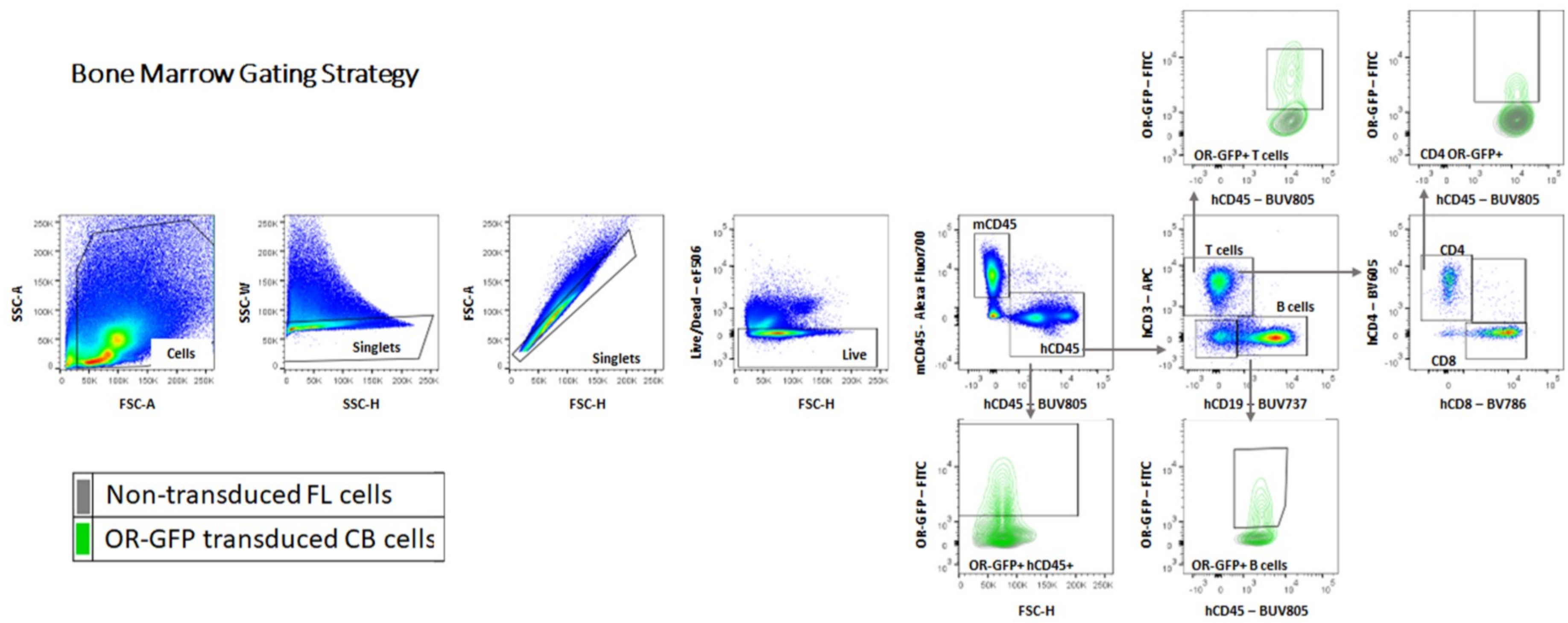


**Fig. S5. Gating strategy for FACS analysis.** Immune cell subsets and representative FACS plots for OR-GFP signal in bone marrow (BM).

C

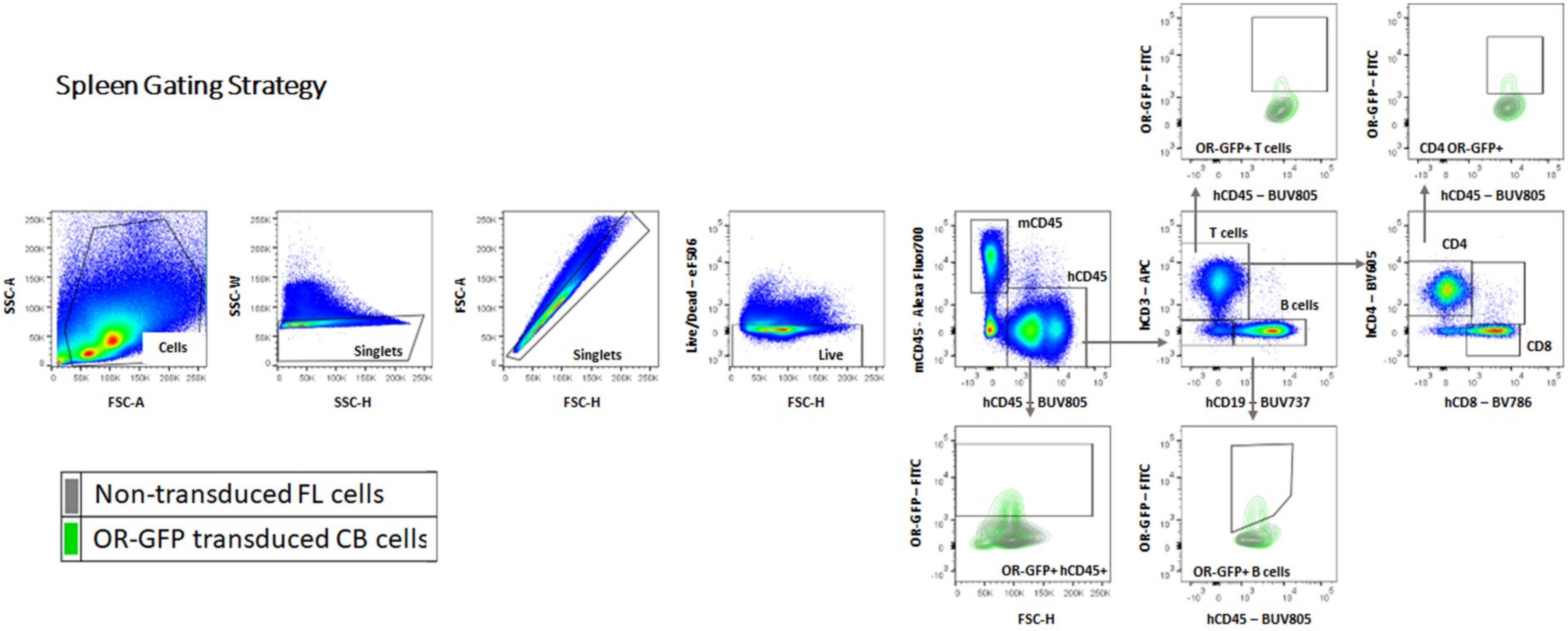


**Fig. S6. Gating strategy for FACS analysis.** Immune cell subsets and representative FACS plots for OR-GFP signal in Spleen.

D

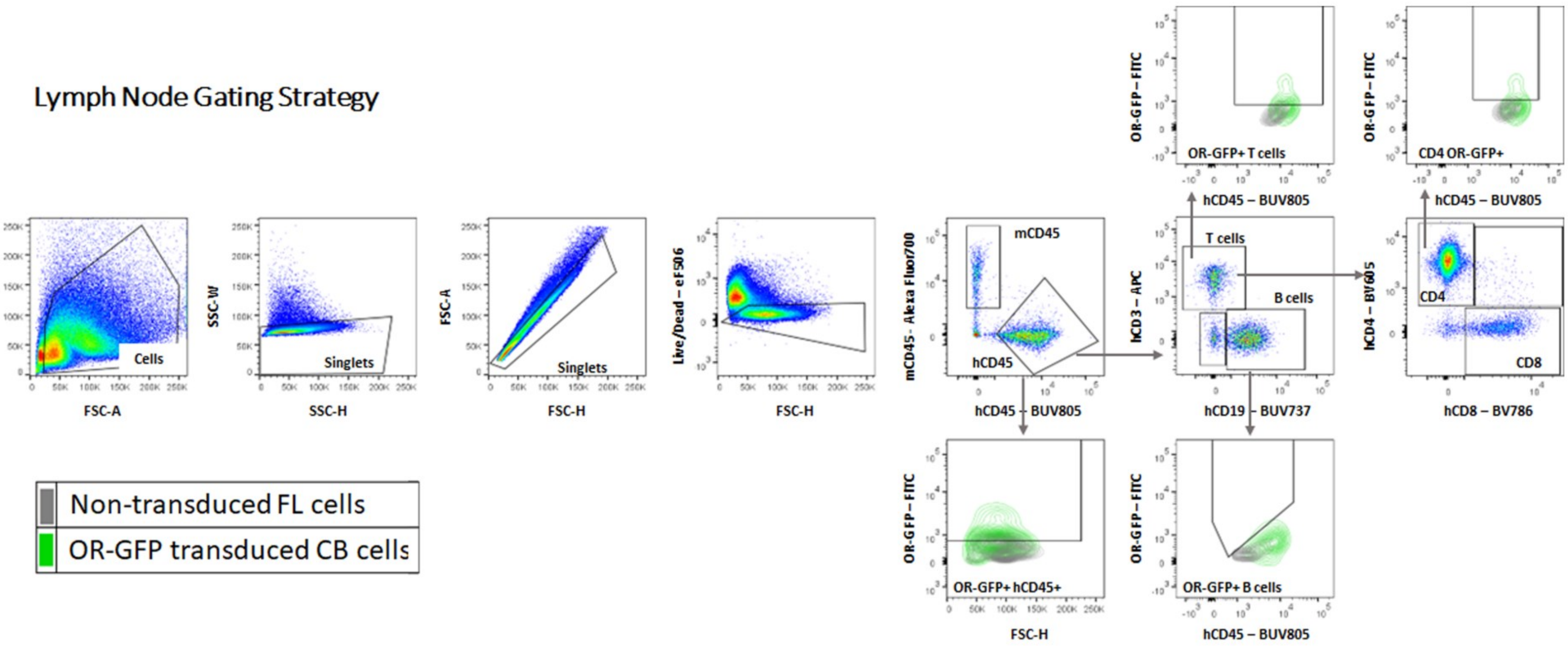


**Fig. S7. Gating strategy for FACS analysis.** Immune cell subsets and representative FACS plots for OR-GFP signal in Lymph nodes (LN).
